# On the interpretation of transcriptome-wide association studies

**DOI:** 10.1101/2021.08.15.456414

**Authors:** Christiaan de Leeuw, Josefin Werme, Jeanne E. Savage, Wouter J. Peyrot, Danielle Posthuma

## Abstract

Transcriptome-wide association studies (TWAS) aim to detect relationships between gene expression and a phenotype, and are commonly used for secondary analysis of genome-wide association study (GWAS) results. Results from TWAS analyses are often interpreted as indicating a genetic relationship between gene expression and a phenotype, but this interpretation is not consistent with the null hypothesis that is evaluated in the traditional TWAS framework. In this study we provide a mathematical outline of this TWAS framework, and elucidate what interpretations are warranted given the null hypothesis it actually tests. We then use both simulations and real data analysis to assess the implications of misinterpreting TWAS results as indicative of a genetic relationship between gene expression and the phenotype. Our simulation results show considerably inflated type 1 error rates for TWAS when interpreted this way, with 41% of significant TWAS associations detected in the real data analysis found to have insufficient statistical evidence to infer such a relationship. This demonstrates that in current implementations, TWAS cannot reliably be used to investigate genetic relationships between gene expression and a phenotype, but that local genetic correlation analysis can serve as a potential alternative.

## Introduction

Transcriptome-wide association studies (TWAS)^1–8^ constitute a statistical framework commonly used to study relationships between gene expression and a phenotype. Combining expression quantitative trait loci (eQTL) and genome-wide association study (GWAS) results, a TWAS analysis estimates the genetic component of the gene expression and then tests whether this is associated with the phenotype. Since TWAS does not require that gene expression and the phenotype are measured in the same sample, it can be used with any phenotype for which GWAS data is available, giving it a broad scope of application.

From a significant TWAS association, it would be tempting to conclude that there is a genetic relationship between the expression of the tested gene and the phenotype of interest, and this interpretation is indeed commonly found in applied TWAS studies^9–13^. Such an interpretation is also suggested by various TWAS method papers such as Gamazon (2015)^7^ and Mancuso (2019)^8^, with for example the latter stating that “[TWAS] can be viewed as a test for non-zero local genetic correlation between expression and trait.” However, the traditional TWAS framework does not directly test the relationship of the phenotype with the genetic component of the gene expression. Rather, it only tests the relationship with the *predicted* genetic component of the gene expression. This is crucial, as traditional TWAS methods do not account for the uncertainty in that prediction^6,7^, meaning they cannot be accurately used to infer a genetic relationship between gene expression and the phenotype on a population level.

In this study, we show that this distinction has important implications for the interpretation and utility of TWAS results. First, we dissect the mathematical structure of the TWAS framework, explicating the null hypothesis that is tested in TWAS analysis, and then demonstrating howthis relates to the genetic relationship between gene expression and the phenotype. Second, we use large-scale simulations to demonstrate that if we were to incorrectly interpret TWAS as testing such genetic relationships, this can result in potentially highly inflated type 1 error rates. Finally, we use a real data application with five well-powered phenotypes to illustrate what these error rate inflations can look like in practice, showing that TWAS cannot be used as a reliable tool for detecting genetic relationships between gene expression and a phenotype of interest.

## Results

### The TWAS framework

The traditional TWAS framework is implemented as a two-stage procedure, as follows^1,6,7^. First, a modelis specified for the expression level *E* of a particular gene 
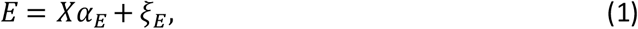
where *X* denotes the genotype matrix of SNPs local to that gene, and *ξ*_*E*_ a residual component independent of *X*. We can further define *G*_*E*_ = *Xα*_*E*_, which reflects the true local genetic component of the gene expression captured by the SNPs in *X*. This model is fitted to a sample with genotype and gene expression data to obtain an estimated genetic effect vector 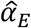. With this, the estimated genetic component of the gene expression is then computed in the GWAS sample for the outcome phenotype *Y* as 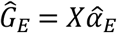. Finally, a linear regression modelof the form 
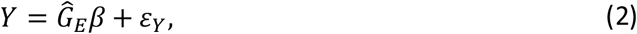
with coefficient *β* and residual *ε*_*Y*_, is used to test the relationship between the estimated genetic component 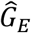 and the phenotype *Y*, evaluating the null hypothesis of no association *H*_0_: *β* = 0.

Note that our presentation here is simplified for the sake of brevity, which includes assuming a continuous phenotype and omitting covariates. The second stage can also be rewritten to require only pre-existing GWAS summary statistics rather than raw genotype and phenotype data as input. See *Methods - Outline of TWAS framework* for a description of the more general case.

In practice, most existing TWAS methods have this mathematical structure^6–8,14–24^, including all the ‘linear’ models listed in Table 1 aside from CoMM^25^. Where they differ is in their implementation, particularly in how 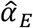 is estimated^6–8,14–25^. Since the number of SNPs in *X* is usually much larger than the sample size of the gene expression data, and are also highly collinear, it is not possible to fit the model in (1) using standard procedures such as Ordinary Least Squares. Most TWAS implementations therefore use a penalized regression approach such as Elastic Net, LASSO or a spike -and-slab model to resolve this issue (see Table 1). Although different kinds of penalized regression procedures will also yield differ different estimates 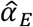, all these methods are still conceptually equivalent, each providing an estimate 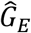 of the genetic component of the gene expression for use in the second stage.

**Table 1.** Overview of available TWAS analysis methods.

| Method | Weight estimation<sup>a</sup> | Base model extensions |
| --- | --- | --- |
| *Linear models* |  |  |
| Gamazon (2015)<sup>7</sup> - PrediXcan | Marginal<br>LASSO<br>Elastic net | - |
| Gusev (2016)<sup>6</sup> - FUSION<sup>b</sup> | Top eQTL<br>BLUP<br>Bayesian LMM<br>LASSO<br>Elastic net | - |
| Mancuso (2017)<sup>14</sup> - RhoGE | BLUP | - |
| Barbeira (2018)<sup>15</sup> - MetaXcan | Marginal<br>LASSO<br>Elastic net | - |
| Su (2018)<sup>16</sup> - MiST | External | Models additional variance component for genetic effects not mediated by predicted expression |
| Hu (2019)<sup>17</sup> - UTMOST | Multivariate LASSO | Simultaneously models multiple tissues during weight estimation |
| Yang (2019)<sup>25</sup> - CoMM | Collaborative mixed model | Estimates weights and associations with phenotype simultaneously in single model |
| Mancuso (2019)<sup>8</sup> - FOCUS | External | Models multiple genes at once, as well as additional pleiotropic genetic effects on phenotype |
| Nagpal (2019)<sup>18</sup> - TIGAR | Dirichlet process regression | Multivariate model with multiple outcome phenotypes |
| Liu (2020)<sup>19</sup> - T-GEN | Spike & Slab | Incorporates epigenetic information into weight estimation process |
| Luningham (2020)<sup>20</sup> - BGW-TWAS | Spike & Slab | Models additional trans-eQTL component |
| Bhattacharya (2021)<sup>21</sup> - MOSTWAS | Elastic net<br>BLUP | Models additional components for trans-eQTL or other molecular phenotypes |
| *Non-linear models* |  |  |
| Xu (2017)<sup>22</sup> - ASPU | External | Uses adaptive test combining sums of powers of score statistics for different powers (includes linear model) |
| Zhang (2020)<sup>23</sup> | External | Uses adaptive test combining linear model with sum of squared score statistics |
| Tang (2021)<sup>24</sup> - VC-TWAS | External | Uses sum of powers of score statistics instead of linear model |

LASSO: least absolute shrinkage and selection operator; BLUP: best linear unbiased predictor; LMM: linear mixed model
<sup>a</sup> Multiple entries for a method denote different options; 'marginal' refers to marginal SNP effect sizes being used as weights, 'external' means the method requires precomputed weights from an external source
<sup>b</sup> The name 'FUSION' and the LASSO and elastic net options for this method were added after publication of the Gusev (2016) paper

Not all existing methods adopt this structure, such as the methods listedas ‘non-linear’ in Table 1, but these methods run into the same issues that will be discussed below for the traditional ‘linear’ TWAS framework (see *Supplemental Information - Non-linear TWAS models* for more details). Throughout this paper, unless otherwise specified, we will use TWAS to refer specifically to the linear TWAS framework as described above.

### Structure of the TWAS null hypothesis

TWAS evaluate the null hypothesis *H*_0_: *β* = 0 for the model presented in equation (2), which be rewritten to mathematically equivalent formulations to provide a clearer understanding of what is being tested. Firstly, we can observe that 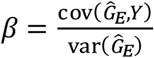, and therefore *β* will be zero if and only if the covariance 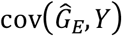 is zero. As such, the TWAS null hypothesis is equivalent to *H*_0_: 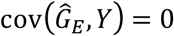.

Secondly, we can partition the phenotype *Y* in the same way as we did with *E* in equation (1), such that

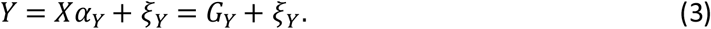

Analogous to *G*_*E*_, this gives us a local genetic component *G*_*Y*_ for the phenotype that is based on the SNPs in *X*, and a residual *ξ*_*Y*_ that is independent of *X* (that is, cov(*X*_*j*_,*ξ*_*Y*_) = 0 for every SNP *j*). Substituting *Y* = *G*_*Y*_ + *ξ*_*Y*_, it follows that 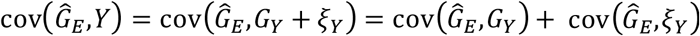, with the last step following from the distributive property of covariance.

Because 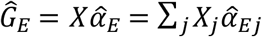 (with *j* indexing the individual SNPs), and as the residual *ξ*_*Y*_ is independentof *X*, it follows that 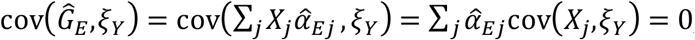, and with that we can conclude that 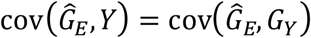. In other words, since *G*_*Y*_ captures all the association that the SNPs in *X* have with the phenotype *Y*, and 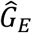is entirely based on *X*, the covariance between 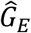and the residual term *ξ*_*Y*_ will be zero by definition. The result of this is that we can further rewrite the TWAS null hypothesis into *H*_0_: 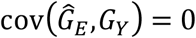.

### Relation to local genetic covariance

Local genetic correlation, analogous to genome-wide genetic correlations, quantifies the genetic relationship between two phenotypes within a particular genomic region, with gene expression, in this context, being one of those phenotypes^26,27^. Since the local genetic correlation can be defined as 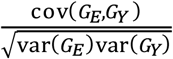, and thus will be zero if and only if cov(*G*_*E*_, *G*_*Y*_) is zero, the genetic relationship between gene expression and the phenotype can be tested using the null hypothesis *H*_0_: cov(*G*_*E*_, *G*_*Y*_) = 0.

However, by contrast, TWAS performs a test of cov 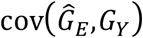rather than cov(*G*_*E*_,*G*_*Y*_), as shown above. To see how these two covariances relate to each other, we can partition the estimated genetic component 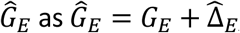, with 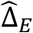the deviation of the sample estimate from the true genetic component *G*_*E*_. Again using the distributive property of covariance, we can then conclude cov 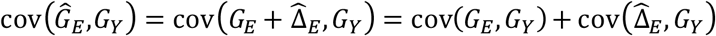. For ease of notation, we will denote 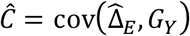. We can thus see that the covariance term 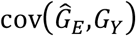 that is tested by TWAS is offset from the local genetic covariance cov(*G*_*E*_,*G*_*Y*_) by *Ĉ*, and plugging this relationship into *H*_0_: 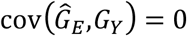, we find that the TWAS null hypothesis is equivalent to *H*_0_: cov *H*_0_ cov(*G*_*E*_,*G*_*Y*_)=− *Ĉ*.

Crucially, the regression model in equation (2) conditions on its predictor 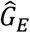, thus considering the value of 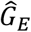 to be fixed. It follows that the values of the deviation term 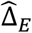, and therefore that of *Ĉ*, are fixed as well, even though 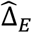 is the result of the random sampling of *E* in the gene expression data. As such, the quantity −*Ĉ* that TWAS implicitly tests cov(*G*_*E*_,*G*_*Y*_) against, will have an unknown, arbitrary value (different for each gene) that reflects the specific error in the estimation that was realized when the gene expression *E* was generated, with the distribution from which this value is drawn also determined by choice of model used for estimating *α*_*E*_ (for more details, see *Supplemental Information – The TWAS null value*).

To further illustrate this, consider an analogy to the t-test. Suppose we are interested in testing whether the population means *μ*_*A*_ and *μ*_*B*_ of some variable for groups *A* and *B* differ. However, instead of performing a two-sample t-test with *H*_0_: *μ*_*A*_ = *μ*_*B*_, or equivalently *H*_0_: *μ*_*A*_ − *μ*_*B*_ = 0, we compute the sample mean 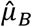 for group *B* and use a single sample t-test with 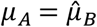. Let us assume that in this case, in our sample from group *B* the realized error in the estimate 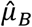 is 1, ie. 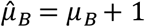. This means that with our single sample t-test, we would be testing *H*_0_: *μ*_*A*_ − *μ*_*B*_ = 1. As with TWAS, because we are disregarding the estimation uncertainty in 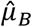 when performing our test, we end up testing the quantity of interest, *μ*_*A*_ − *μ*_*B*_, against an unknown null value, which isn’t very informative.

### Interpretation of TWAS results

As shown above, TWAS implicitly tests the local genetic covariance cov(*G*_*E*_,*G*_*Y*_) against −*Ĉ* (for an unknown value of *Ĉ*) rather than zero, and as such it cannot be considered a direct test of the genetic relationship between gene expression and the phenotype. This raises the question of how TWAS results actually *can* be interpreted, which we can address by determining the circumstances in which the TWAS null hypothesis *H*_0_: 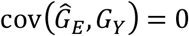 is true.

As shown above, the null hypothesis *H*_0_: 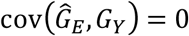 is equivalent to *H*_0_: cov(*G*_*E*_,*G*_*Y*_) = −*Ĉ*. Examining the structure of *Ĉ* more closely we can see that it decomposes into a weighted sum 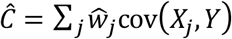 of the (population-level) covariances cov(*X*_*j*_, *Y*) of the SNPs in *X* with the phenotype (see *Supplemental Information – The TWAS null value*). These weights 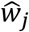 reflect the error in the effect size estimate for each SNP *j*, ie. 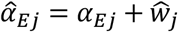, and as such their values are the realization of a random process.

It follows that, if there is any genetic association between the SNPs in *X* and the phenotype *Y*, the TWAS null hypothesis will only be true if by chance the values of 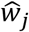 are such that 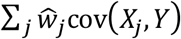 happens to equal −cov(*G*_*E*_, *G*_*Y*_). Because the value of 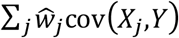 is effectively a random draw from a continuous space, the probability of this happening is infinitessimal. Realistically, the TWAS null hypothesis will therefore only be true if there is no genetic association between *X* and *Y*, with cov(*X*_*j*_,*Y*) = 0 for every SNP *j*. In this case, 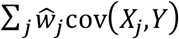 will be zero as well, and at the same time the absence of such genetic signal also means that the variance of *G*_*Y*_ is zero, and therefore no covariance with *G*_*E*_ can exist. As such, in this case both cov(*G*_*E*_,*G*_*Y*_) and *Ĉ* will be zero, and *H*_0_: cov(*G*_*E*_, *G*_*Y*_) = −*Ĉ* is therefore true.

In practice, the TWAS null hypothesis will therefore be true if, and only if, the SNPs in *X* are all independent of the phenotype, and a test of this null hypothesis therefore amounts to a joint genetic association test, ie. a test of independence between *X* and *Y* akin to testing *H*_0_: *α*_*Y*_ = 0 in the general regression model in equation (3), though with somewhat different power characteristics. Although as shown, the TWAS null hypothesis *H*_0_: cov(*G*_*E*_,*G*_*Y*_) = −*Ĉ* being true does imply that the null hypothesis *H*_0_: cov(*G*_*E*_, *G*_*Y*_) = 0 of no local genetic relationship is true as well, the reverse does not hold. If genetic association does exist between *X* and *Y*, then *H*_0_: cov(*G*_*E*_,*G*_*Y*_) = −*Ĉ* will be false, but as long as this genetic association is independent of the genetic signal captured by *G*_*E*_, then cov(*G*_*E*_,*G*_*Y*_) will still be zero. This is why it does not follow from the TWAS null hypothesis being false that a local genetic relationship exists between gene the expression and the phenotype.

Despite this, the intuition may remain that even though 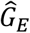 is uncertain, it is still based on the true *G*_*E*_ and should therefore still contain *some* information relevant to the genetic relationship between gene expression and the phenotype. And this is indeed the case, because as shown the tested covariance 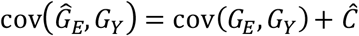 is partly based on the true local genetic covariance cov(*G*_*E*_, *G*_*Y*_). But although present, this information cannot meaningfully be extracted from the TWAS results, since even when 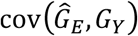 is found to deviate significantly from zero, the TWAS cannot tell us whether this is due to a strong local genetic relationship through cov(*G*_*E*_,*G*_*Y*_), or just through the noise term *Ĉ*.

### TWAS as a test of genetic relationship

As established above, TWAS does not technically provide a direct test of the genetic relationship between gene expression and the phenotype, since its null hypothesis is equivalent to *H*_0_: cov(*G*_*E*_, *G*_*Y*_) = −*Ĉ*, with the value of *Ĉ* unknown. However, it is possible that in the scenarios that are likely to arise in practice, the value of *Ĉ* might be so small as to be negligible. If so, we could simply treat TWAS as if it is testing the null hypothesis *H*_0_: cov(*G*_*E*_,*G*_*Y*_) = 0 after all, and interpret its significant results as indicative of genetic relationships between gene expression and the phenotype without issue.

In the remainder of this paper, we will evaluate the viability in practice of treating TWAS as a test of the null hypothesis *H*_0_: cov(*G*_*E*_,*G*_*Y*_) = 0, first using large-scale simulations to determine the resulting type 1 error rates across a range of different scenarios, and then using an application to real data to illustrate the impact in practice. We used FUSION^6^ for this evaluation, as it is a widely used tool, and it implements a range of different models within the same tool that are representative of those used in many other TWAS methods (see Table 1). We also included CoMM^25^, since it uses a setup that is somewhat different than that of the other TWAS methods, and closer to that of local genetic correlation methods^26,27^ (see *Supplemental Information - Comparison with the CoMM model*). Note that reported type 1 error rates will be relative to *H*_0_: cov(*G*_*E*_,*G*_*Y*_) = 0, rather than the null hypotheses these methods are designed to test, as the aim is to evaluate the viability of using these methods when they are treated as a test the genetic relationship between gene expression and the phenotype (as they are often interpreted in practice).

To serve as a reference, we included the local genetic correlation model implemented in LAVA (denoted LAVA-rG), which is designed and calibrated to test the null hypothesis *H*_0_:cov(*G*_*E*_,*G*_*Y*_) = 0 directly. Furthermore, we also implemented a TWAS model in the LAVA framework (denoted LAVA-TWAS), which is identical to LAVA-rG in every way, except that like other traditional TWAS methods it treats 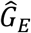 as given (rather than modelling its distribution under sampling of the gene expression *E*, like LAVA-rG does; see *Methods - LAVA implementation of TWAS*). This implementation allows for a direct comparison between testing *H*_0_: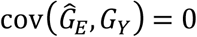 and testing *H*_0_: cov(*G*_*E*_, *G*_*Y*_) = 0, unconfoundedby other differences in model or implementation.

### Simulation study

To evaluate the performance of TWAS when interpreted as a test of the genetic relationship between gene expression and outcome phenotype, we performed a series of simulations, generating gene expression and phenotype values under the null hypothesis *H*_0_: cov(*G*_*E*_,*G*_*Y*_) = 0 and computed type 1 error rates. Genotype data from the UK Biobank^28,29^ was used as a basis for the simulations, and we varied the number of SNPs per simulated gene, as well as the sample size and local heritability of the gene expression and outcome phenotype data (see *Methods - Simulation study*). Analysis was performed using LAVA-TWAS and LAVA-rG, as the four different models in FUSION (BLUP, Bayesian LMM, Elastic Net, LASSO), and CoMM.

The simulations showed considerable inflation of type 1 error rates (*α* = 0.05) when using LAVA-TWAS to evaluate *H*_0_: cov(*G*_*E*_, *G*_*Y*_) = 0, depending on the simulation condition. The inflation strongly increased with larger local heritability or sample size for the outcome phenotype (Figure 1), with the local heritability also interacting with the number of SNPs in the gene (Supplemental Figure 1). In addition, some decrease in the type 1 error rate inflation was also found with greater local heritability or sample size for the gene expression, but this effect is much less pronounced. Moreover, the type 1 error rate inflation also became progressively worse when evaluating it at stricter significance thresholds than the *α* of 0.05 used so far (Figure 2). Type 1 error rates also increased when filtering out genes that exhibit insufficient genetic signal for the gene expression, as is general practice in TWAS analysis (Supplemental Figure 2). To validate the LAVA-TWAS model, type 1 error rates for LAVA-TWAS under its actual null hypothesis of *H*_0_: 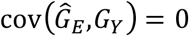 were also assessed. These were found to be well controlled in all conditions (Supplemental Figure 3), further emphasizing the disconnect between this null hypothesis and the null hypothesis of no genetic relationship.

**Figure 1.**
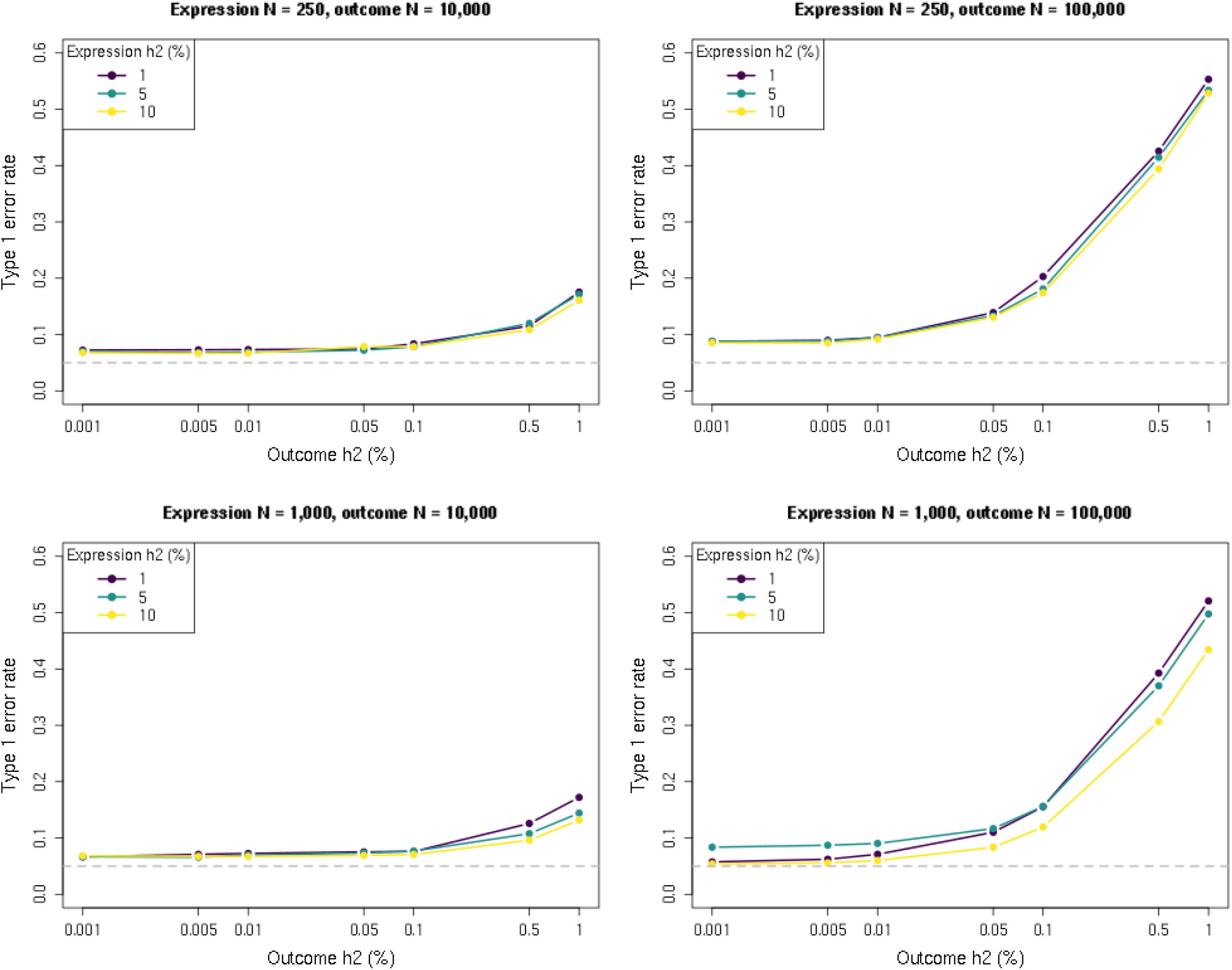
Type 1 error rate simulation results as a function of sample size and local heritability. Shown is the type 1 error rate (at significance threshold of 0.05) of the LAVA-TWAS model for the null hypothesis of no genetic covariance (cov(G_E_, G_Y_) = 0), at different levels of local heritability (h2) for outcome phenotype (horizontal axis) and gene expression (separate lines). Results are shown for different sample sizes (N) for the expression (rows) and outcome (columns) data, and at 1,000 SNPs. As shown, the type 1 error rates get considerably larger with increasing sample size or local heritability for the outcome. Conversely, increasing these for the expression reduced the type 1 error rate, but the effect of this is much less pronounced.

**Figure 2.**
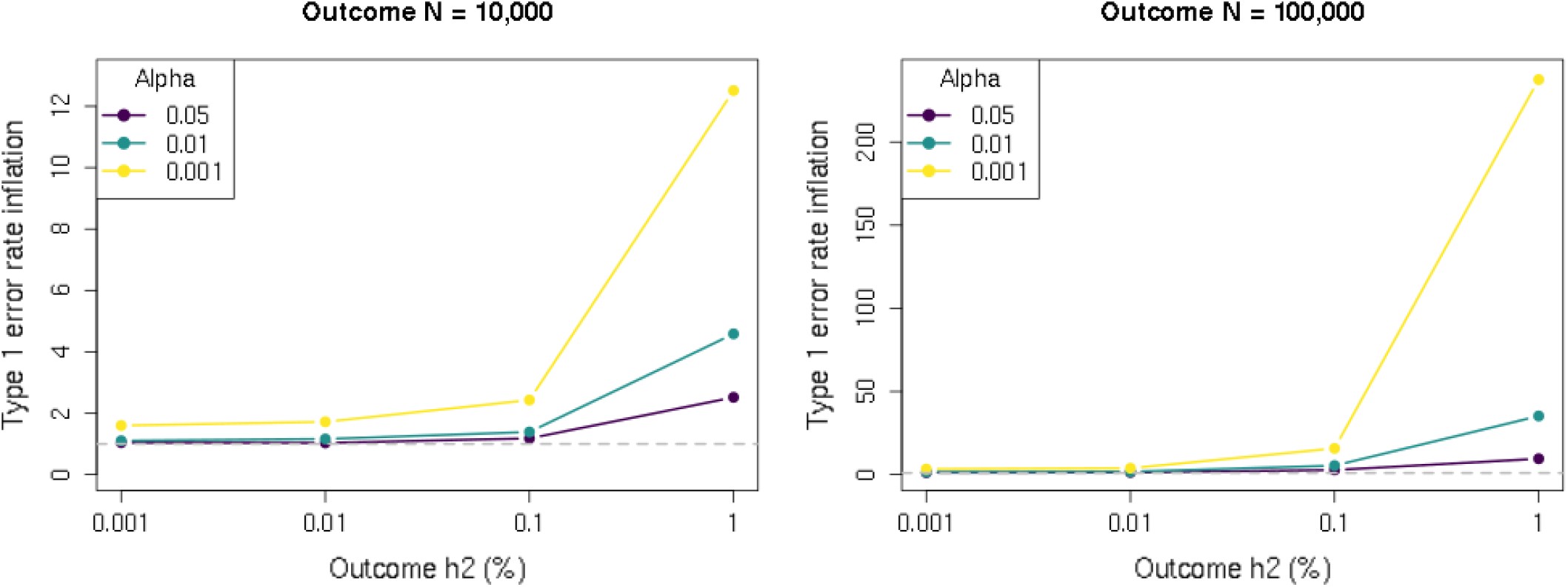
Type 1 error rate inflation simulations results as a function of significance threshold. Shown is the type 1 error rate inflation of the LAVA-TWAS model for the null hypothesis of no genetic covariance (cov(G_E_, G_Y_) = 0), for different levels of significance threshold α. The type 1 error rate inflation is defined as the type 1 error rate divided by the significance rate α, and equals 1 if the error rates are well-controlled. Results are shown for simulations with expression sample size of 1,000, expression local heritability of 5%, and 1,000 SNPs. As shown, the type 1 error rate becomes relatively more inflated the lower the significance threshold used (separate lines), with the difference becoming more pronounced at higher local heritability (h2) (horizontal axis) and sample size (N) (separate panels) for the outcome.

Results for the four FUSION penalized regression models are largely the same as for LAVA-TWAS (Figure 3), suggesting that the type of model and penalization usedfor the estimation of 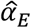 does not strongly affect the subsequent type 1 error rate inflation. Although the Elastic Net and LASSO models showed lower inflation than other models in the 1% gene expression heritability conditions, this can be explained by the fact that they frequently failed to converge in these conditions when estimating equation (1). This difference was no longer present in the 5% heritability conditions (see Supplemental Figure 4).

**Figure 3.**
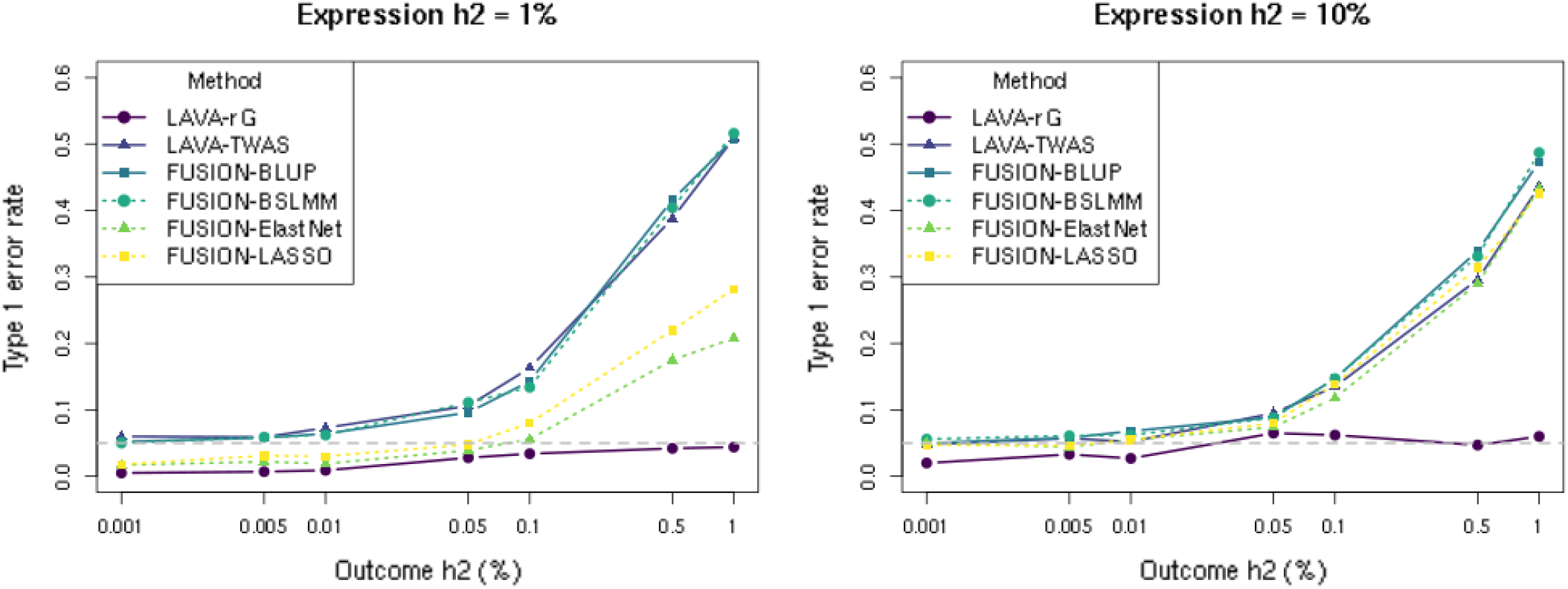
Comparison of type 1 error rate simulation results across models. Shown is the type 1 error rate (at significance threshold of 0.05) of the five different TWAS models and LAVA-rG for the null hypothesis of no genetic covariance (cov(G_E_, G_Y_) = 0), at different levels of local heritability (h2) for outcome phenotype (horizontal axis) and gene expression (separate panels). Results are shown for sample sizes of 1,000 and 100,000 for the expression and outcome respectively, with 1,000 SNPs. As shown, the results are very similar across the different models. The lower type 1 error rate for Elastic Net and LASSO compared to the other TWAS models for the 1% expression heritability condition is due to these two models sometimes failing to converge at very low genetic signal for the expression (see also Supplemental Figure 4).

Similar patterns of type 1 error rate inflation were found for CoMM as well and were genera **l**y more pronounced, though the influence of the gene expression heritability was more complex than for the other methods (Supplemental Figure 5). This is due to specific constraints in the model, which are analogous to the 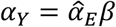 constraint the traditional TWAS models impose the multiple regression in equation (4) (see *Supplemental Information - Comparison with the CoMM model*). As in the validation simulations in Werme (2022)^26^, type 1 error rates for LAVA-rG were found to be well-controlled (Figure 3).

Overall, the results of our simulations demonstrate that treating TWAS analysis as if they were a test of the null hypothesis of no genetic relationship *H*_0_: cov(*G*_*E*_, *G*_*Y*_) = 0 can cause considerable inflation of the type 1 error rates in many conditions. A strong effect was observed for the sample size and heritability of the outcome phenotypes, in line with the interpretation of TWAS as a joint association test. Of particular concern is that the inflation increases when filtering on the univariate signal of the gene expression, which is commonly done in TWAS studies. The inflation also increases when using lower significance thresholds, which is likely to apply in practice to correct for multiple testing, and often at more stringent levels than shown in our simulations. See *Supplemental Information - Discussion of simulation results* for more discussion of the findings from these simulations.

### Real data analysis

To gauge the magnitude of the impact this issue has in a real data context, we applied TWAS to GWAS results of five well-powered phenotypes^30–33^ (see Table 2), with eQTL data for blood gene expression from GTEx^34^ (v8). A total of 14,584 genes were analysed, performing TWAS analysis for those genes for which the univariate gene expression signal was significant at *α*_BONF_ = 0.05/14,584 = 3.43 × 10^−6^ (see *Methods-Data* and *Methods - Realdata analysis*). The data was analysed with the Elastic Net and LASSO models in FUSION and LAVA-TWAS. We analyzed the data with LAVA-rG as well to provide a baseline for the LAVA-TWAS results, and determine for how many associations the *H*_0_: cov(*G*_*E*_, *G*_*Y*_) = 0 null hypothesis can be validly rejected.

**Table 2.** Summary of results of TWAS and local genetic correlation analyses of published summary statistics for five phenotypes.

| Phenotype | Sample size<sup>a</sup> | Number of SNPs<sup>b</sup> | Genes tested | Significance threshold | LAVA signif. associations |  |  | FUSION signif. associations |  |
| --- | --- | --- | --- | --- | --- | --- | --- | --- | --- |
| | | | | | $r_G$ | TWAS | Overlap %<sup>c</sup> | Elast. net | LASSO |
| Blood pressure<sup>33</sup> | 361K | 5.94M | 6,686 | $7.48 \times 10^{-6}$ | 68 | 107 | 63.6 | 100 | 109 |
| BMI<sup>30</sup> | 807K | 6.28M | 5,453 | $9.17 \times 10^{-6}$ | 134 | 232 | 57.8 | 297 | 328 |
| Type 2 diabetes<sup>33</sup> | 18.5K/366K | 5.94M | 6,686 | $7.48 \times 10^{-6}$ | 27 | 45 | 60.0 | 26 | 33 |
| Educational attainment<sup>31</sup> | 766K | 6.18M | 5,532 | $9.04 \times 10^{-6}$ | 59 | 108 | 54.6 | 149 | 165 |
| Schizophrenia<sup>32</sup> | 67.4K/94.0K | 6.08M | 5,765 | $8.67 \times 10^{-6}$ | 53 | 92 | 57.6 | 109 | 114 |

<sup>a</sup> Showing case/control for binary phenotypes
<sup>b</sup> After filtering for overlap with 1,000 Genomes and GTEx SNPs
<sup>c</sup> Percentage of significant LAVA-TWAS associations that were also LAVA-rG associations

A summary of the results is given in Table 2 and as shown, the number of significant LAVA-TWAS associations is considerably higher than those for LAVA-rG, with on average only about 59% of LAVA-TWAS associations confirmed by LAVA-rG. From this it follows that if we were to interpret LAVA-TWAS as a test of genetic relationship between gene expression and the phenotype, 41% of the associations found would not be valid. This is because LAVA-rG is calibrated for testing *H*_0_: cov(*G*_*E*_, *G*_*Y*_) = 0, and LAVA-TWAS only differs from LAVA-rG in the fact that it conditions on 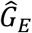, whereas LAVA-rG models the distribution of the estimation uncertaintyin 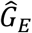that we know exists. This single difference is the cause of the inflated type 1 error rates observed for LAVA-TWAS (but not LAVA-rG) in the simulations. Consequently, because there is no other difference between the two, the 41% of significant LAVA-TWAS associations not shared with LAVA-rG cannot be due to a legitimate increase in power, meaning that they are not valid if interpreting LAVA-TWAS as a test of *H*_0_: cov(*G*_*E*_, *G*_*Y*_) = 0 (see also *Supplemental Information – Underlying distributions*).

Although no equivalent reference model directly testing *H*_0_: cov(*G*_*E*_,*G*_*Y*_) = 0 is available for the FUSION models, the FUSION results paint a similar picture. The number of associations found when using FUSION are at a comparable level to LAVA-TWAS (Table 2, Supplemental Table 1), and the FUSION models had very similar type 1 error rate inflation levels as LAVA-TWAS in the simulations as well. Taken together, it can reasonably be concluded that the proportion of FUSION results in the real data analysis that are not supported by sufficient statistical evidence, when interpreted as a test of the genetic relationship between gene expression and the phenotype, is likely at a very similar level as LAVA-TWAS

We also evaluated results for genes with a univariate p-value for the gene expression greater than 0.05. As shown in Supplemental Table 2, virtually no local genetic correlations are found with LAVA-rG for these genes, yet results for the TWAS analyses remain at a similar level as in the primary analysis in Table 2. Because these genes have no detectable genetic component for the gene expression, they would typically not be included in a TWAS analysis. But the fact that TWAS still yields a considerable number of significant associations in this context, when 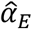 reflects essentially just noise, serves to further illustrate how TWAS simply functions as a form of joint association test, with results not necessarily reflecting any genetic relationship between gene expression and the outcome phenotype.

### Deflation of standard errors

Fundamentally, the statistical reason behind the results in our simulations and real data analysis is that the TWAS model omits a source of variance that we know is present. The estimated genetic effect vector 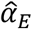 is subject to estimation uncertainty due to the sampling of the gene expression *E*, but this uncertainty is not accounted for in the TWAS test. If the aim is to evaluate the genetic relationship with the phenotype, by testing the null hypothesis *H*_0_: cov(*G*_*E*_,*G*_*Y*_) = 0, the resulting standard errors will therefore only reflect the uncertainty due to sampling of *Y*, and not that of *E*. As such, the standard errors will be too small, creating a downward bias in the p-values (see *Supplemental Information – Underlying distributions* for more details). For LAVA-TWAS the corrected standard errors are known, as these are the LAVA-rG standard errors, and we can therefore use these to illustrate the issue. We computed this standard error deflation individually for each gene in our analysis, as the ratio of the LAVA-TWAS standard errors to their corrected value. These are summarized in Table 3 and, as shown, they vary considerably across genes, with negligible deflation for some genes but severe deflation for others. Figure 4 illustrates the relation between this standard error deflation and the resulting bias in p-values, using the BMI results as an example, with Supplemental Figure 6 further showing the level of realized bias. This high variability of standard error deflation across genes also demonstrates that this issue must be addressed at the level of the individual gene, and cannot be resolved by applying a simple post-hoc correction to the gene p-values.

**Table 3.** Summary of TWAS standard error deflation values per gene.

| Phenotype | Mean | Minimum | Quantiles |  |  |  |  |
| --- | --- | --- | --- | --- | --- | --- | --- |
|  |  |  | 5% | 25% | Median | 75% | 95% |
| Blood pressure | 0.93 | 0.69 | 0.84 | 0.90 | 0.94 | 0.97 | 0.99 |
| BMI | 0.88 | 0.46 | 0.75 | 0.84 | 0.89 | 0.94 | 0.98 |
| Type 2 diabetes | 0.95 | 0.51 | 0.89 | 0.93 | 0.96 | 0.98 | 1.00 |
| Educational attainment | 0.93 | 0.62 | 0.84 | 0.90 | 0.94 | 0.97 | 0.99 |
| Schizophrenia | 0.92 | 0.65 | 0.83 | 0.89 | 0.92 | 0.96 | 0.99 |

**Figure 4.**
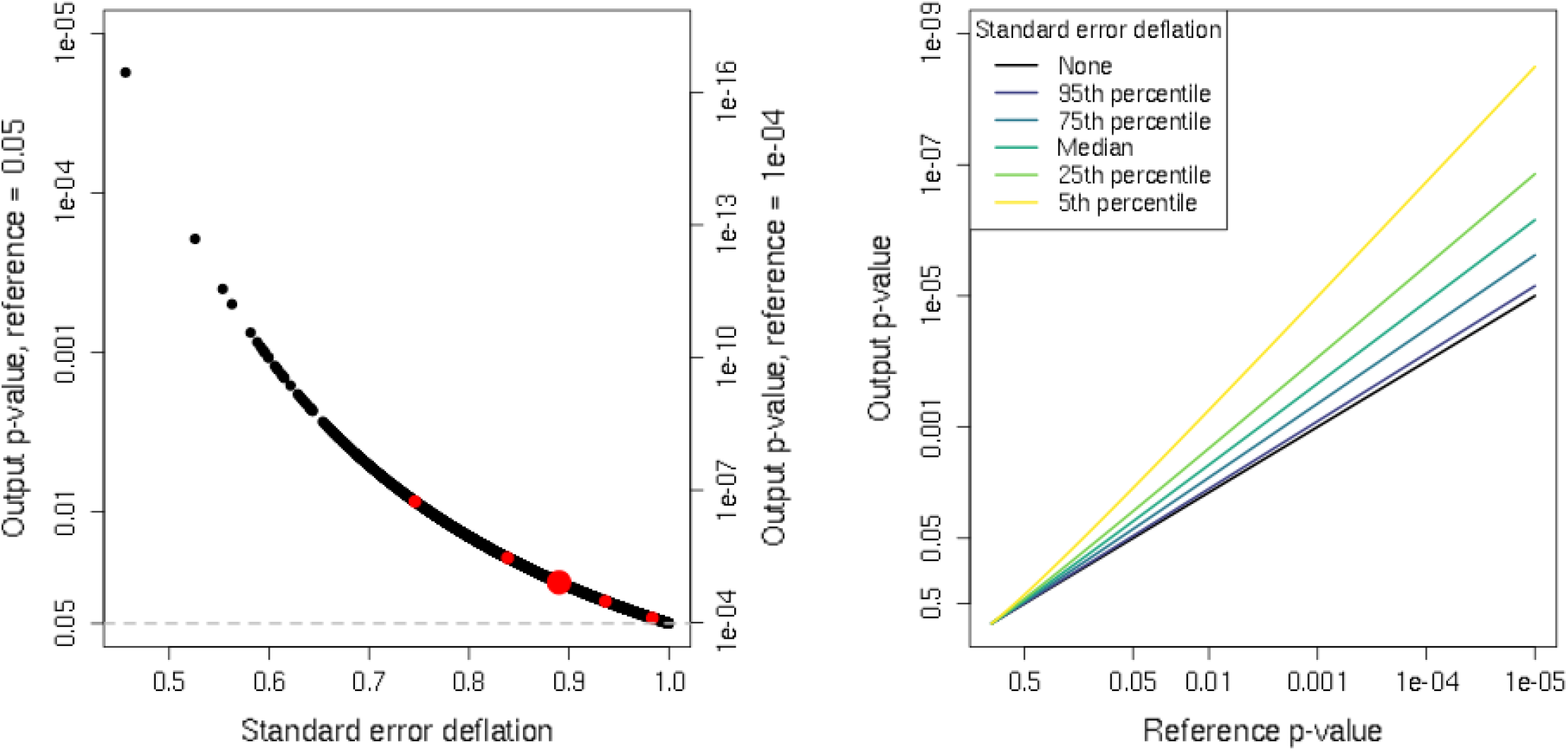
Illustration of p-value bias as a result of deflated standard errors. Shown in the left panel are the p-values that will result from a LAVA-TWAS analysis at different levels of standard error deflation, when the true p-value computed with correct standard errors would either be 0.05 (left vertical axis) or 0.0001 (right vertical axis). The standard error deflation values are taken from the BMI analysis, with the large red point denoting the median and the other red points the 5^th^, 25^th^, 75^th^ and 95^th^ quantiles of their distribution (see Table 3). As shown, the relative bias is stronger the lower the p-value. For example, at the median standard error deflation of 0.89, at a reference p-value of 0.05 the biased p-value is about 0.028, about a 1.8-fold decrease in the p-value relative to its true value. At a reference p-value of 0.0001 however, the biased p-value is about 1.2 × 10^−5^, an 8.1-fold decrease. In the right panel is the general relationship between the true reference p-value and the p-value that would result at different levels of standard error deflation, based on the percentiles for the BMI analysis. This further illustrates the relative increase in bias for lower p-values, with the gap with the reference p-value (black line) getting larger at lower reference p-values. Note that all axes with p-values use a -log10(p-value) scaling, but labels are expressed as p-values for ease of interpretation.

## Discussion

In this paper, our aim was to elucidate the interpretation of the TWAS null hypothesis and results. Although TWAS results are often interpreted as indicating a genetic relationship between gene expression and the phenotype, we showed that this is not what the TWAS null hypothesis allows us to test. Instead, the null hypothesis that the traditional TWAS models actually evaluate effectively makes them a test of joint genetic association between the SNPs included in the analysis and the phenotype, irrespective of any genetic relationship with the gene expression. However, because it is not uncommon for TWAS results to be misinterpreted as a test of such a genetic relationship, we aimed to determine to what extent this interpretation was viable in practice. To this end, we performed simulations to determine the degree of type 1 error rate inflation that occurs when treating TWAS results as a test of genetic relationship, and performed real data analysis to demonstrate what this translated to in practice.

Our results showed that using TWAS to infer genetic relationships between gene expression and the phenotype can lead to strongly inflated type 1 error rates, as well as a substantial proportion of invalid significant results, for which such a genetic relationship is insufficiently supported by the statistical evidence in the data. This was further emphasized by the fact that in our real data analysis, TWAS still yielded a considerable number of significant associations even when genetic signal for the gene expression was absent altogether, clearly illustrating how TWAS functions simply as a form of joint association test. The issue is further complicated by the fact that the degree to which standard errors are deflated varies across genes, depending on factors such as the gene size and level of genetic signal for both gene expression and phenotype. This therefore rules out the possibility of a simple post-hoc correction on existing TWAS results, making them difficult to interpret and use in practice. Although TWAS results can still be validly interpreted as a form of joint association test for the genetic relationship between the SNPs and the phenotype, as out results show, it does not allow for any conclusions to be drawn about the relationship of the phenotype with the gene expression, and therefore does not address the research questions that most researchers are looking to answer with such data. While the issue of uncertainty in eQTL estimates has occasionally been mentioned in TWAS literature^1,25,35,36^, its implications for the validity of the TWAS framework have received little scrutiny thus far. Moreover, although various TWAS methods have included simulations to evaluate type 1 error rates, these all either treated 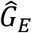 as fixed in the simulations just as in the TWAS model itself^16,23^, or only simulated under a null scenario of the phenotype having no genetic component at all (ie. *G*_*Y*_ = 0)^22,24,25^. This explains why the issue demonstrated in this paper has not yet been widely discussed.

Addressing this issue with TWAS generally requires fully accounting for the uncertainty of the estimate 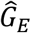 in relation to the true genetic component *G*_*E*_. While it is in principle possible to incorporate this into existing TWAS models, in practice, the feasibility of implementing this will strongly depend on the approach used to fit the model in equation (1), as the distribution of the noise component induced by penalized regression models can take mathematically very complex forms.

Another option is to use alternative methods to detect genetic relationships between gene expression and a phenotype. As demonstrated, local genetic correlation analysis methods could potentially fill this role, as these are explicitly designed to evaluate local genetic relationships between phenotypes. However, methods like colocalization^37,38^ or Mendelian Randomization (MR)^39–41^ could potentially be used for this as well. These methods use quite different model assumptions, but test null hypotheses that can answer similar research questions pertaining to local genetic relationships between gene expression and phenotypes. Of note as well is that the issue with TWAS described in this paper has a clear analogy to the issue of the “no measurement error” (NOME) assumption in MR analysis^42^. As such, strategies used to remove this NOME assumption in the MR context may also help inform solutions for addressing the issue in TWAS methods.

Investigating genetic relationships between gene expression and phenotypes, as TWAS is often intended to do, has the potential to provide valuable insights into the genetic etiology of many phenotypes. But although traditional TWAS models are statistically valid relative to their own null hypothesis, ultimately they only perform a test of joint association of the phenotype with the SNPs in a region, and thus cannot be used to reliably draw conclusions about any genetic relationship with gene expression. Further development of existing TWAS methods to account for this issue, or adoption of alternative methods such as local genetic correlation analysis, will therefore be essential to support future investigations of the relationship between gene expression and phenotypes.

## Methods

### Ethics statement

This study relied on simulated data based on the UK Biobank genotype data^28^, as well as secondary analysis of publicly available summary statistics. Ethical approval for this data was obtained by the primary researchers.

### Outline of TWAS framework

For a standardized genotype matrix *X* containing SNPs for a particular gene, and with *E* and *Y* the centered gene expression for that gene and outcome phenotype respectively, the general TWAS framework consists of a two-stage procedure, based on the equations 
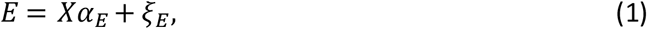
and 
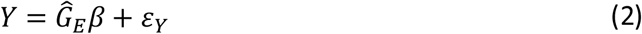
with the estimated genetic component 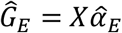, also given in the main text. For ease of notation, we have omitted model intercepts and covariates from these equations, but in practice these will usually be included. An alternative model may also be used rather than the linear regression in equation (2), such as a logistic regression if the phenotype is binary.

For each gene and tissue, equation (1) is first fitted to the eQTL data to obtain the estimated weight vector 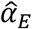. This is then used to compute 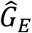 in the target GWAS sample in the second stage, and plugged into the linear model in equation (2). A p-value is then obtained by performing a test on the coefficient *β*. Note that usually the second stage is only performed for genes and tissues that exhibit sufficient genetic association in the eQTL data. This second stage can also be rewritten in terms of GWAS summary statistics, allowing TWAS to be performed without having direct access to the GWAS sample. In this case a genotype matrix *X* obtainedfrom a separate reference sample is used to estimate LD.

Which SNPs are included in *X* varies, but a common choice is to use all available SNPs within one megabase of the transcription region of the gene. Although for simplicity the same genotype matrix *X* is used for equation (1) and (2), there will be separate *X* genotype matrices for each sample. The analysis is therefore restricted to using only those SNPs that are available in both samples, as well as in the LD reference sample when using summary statistics as input.

In practice, equation (1) cannot be fitted with a traditional multiple linear regression model, due to the high LD between SNPs (leading to extreme collinearity), and the number of SNPs typica **l**y exceeding the sample sizes of eQTL data. Some form of regularization in the regression model is therefore required to obtain 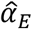, and consequently one of the main discrepancies between TWAS implementations is the specific regularization used (see Table 1). In some cases, rather than fitting equation (1), the elements of 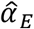 are simply set to the marginal SNP effect estimates instead.

Note that some methods^22–24^ diverge from this linear model structure (Table 1, *non-linear models*). Statistically these can be seen as generalizations of the TWAS framework, though conceptually they can no longer be interpreted as imputing the genetic component of gene expression. See *Supplemental Information - Non-linear TWAS models* for more details.

### Local genetic correlation

The LAVA implementation of local genetic correlation analysis has been described in detail in Werme et al. (2022)^26^. In brief, LAVA uses summary statistics and a reference genotype sample to fit equations (1) and (3), obtaining estimates of 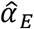 and 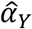 as well as a corresponding sampling covariance matrix for each (a logistic regression equivalent is used for binary phenotypes). To do so, a singular value decomposition for *X* is computed, pruning away excess principal components to attain regularization of the models and allowing them to be fitted.

With the pruned and standardized principal component matrix *W* = *XR* (with *R* the transformation matrix projecting the genotypes onto the principal components), we can write *G*_*E*_ = *Wδ*_*E*_ and *G*_*Y*_ = *Wδ*_*Y*_, where *δ*_*E*_ and *δ*_*Y*_ are the genetic effect size vectors for these principal components. Their estimates 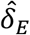 and 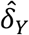 can be used to obtain 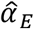 and 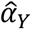 by reversing the transformation through *R*, such that 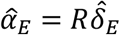 and 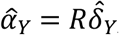, with the projection to the principal components effectively providing a form of regularization in the estimation of *α*_*E*_ and *α*_*Y*_. In practice however, LAVA is defined and implemented directly in terms of *δ*_*E*_ and *δ*_*Y*_ and its estimates, rather than working with *α*_*E*_ and *α*_*Y*_ explicitly.

For ease of notation, we define the combined matrix *G* = (*G*_*E*_, *G*_*Y*_) = *Wδ* for the genetic components, with combined effect size matrix *δ* = (*δ*_*E*_, *δ*_*Y*_). We denote the effect sizes for a single principal component *j* as *δ*_*j*_, corrresponding to the *j*th row of *γ*, and denote the number of principal components as *K*.

The estimates 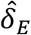 and 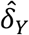 are obtained by reconstructing multiple linear regressions from the input summary statistics (see Werme et al. (2022)^26^ for details). This uses two separate equations of the form *E* = *Wδ*_*E*_ + *ζ*_*E*_ and *Y* = *Wδ*_*Y*_ + *ζ*_*Y*_, analogous to equations (1) and (3) but regressing on *W* rather than *X*, with residual variances 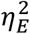 and 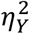 for *ζ*_*E*_ and *ζ*_*Y*_. From these models we have estimates of the form 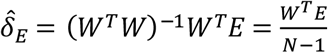 (since *W*^*T*^*W* = *I* (*N* − 1), with *I*_*K*_ the size *K* identity matrix) and similarly 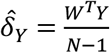, with corresponding sampling distributions 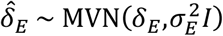 and 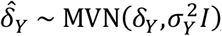, where 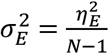 and 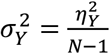 are the sampling variances (ie. squared standard errors). In practice, we obtain these by estimating 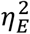 and 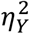 and plugging these in to get estimates 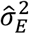 and 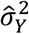.

For principal component *j* we therefore have 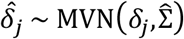, where the diagonal elements of 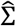 are 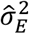 and 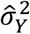 and the off-diagonal elements are 0 (in the general case the off-diagonal elements represent the sampling covariance resulting from sample overlap, but this is not present in the analyses in this study). Since for the covariance matrix of *G*, denoted Ω, we have 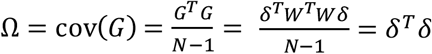, it follows that inference on Ω can be performed using the sampling distributions for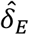 and 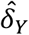 directly.

Using this model, separate univariate tests of joint association of the SNPs in *X* with *E* and *Y* can be performed, testing the null hypotheses 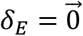 and 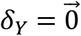 respectively (using standard linear regression F-test for continuous phenotypes (such as gene expression), or a χ^2^ test for binary phenotypes). This is equivalent to testing the local genetic variances 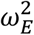 and 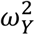, the diagonal elements of Ω. In general, it is recommended in LAVA to test both genetic variances before performing the bivariate analysis, since genetic covariance can only exist in a genomic region where there both phenotypes exhibit some degree of genetic variance, though analogous to standard practice in TWAS literature in this paper we only test on the genetic variance for the gene expression.

From the above distributions it follows that the expected value 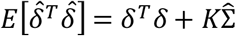, and we can therefore use the method of moments to estimate Ω as 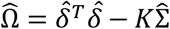. Since in the present analyses there is assumed to be no sample overlap, the off-diagonal elements of Σ are 0, and the estimate for the genetic covariance therefore reduces to 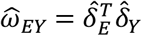. The matrix 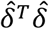has a non-central Wishart sampling distribution, which in the current implementation of LAVA is used to obtain p-values to test *H*_0_: *ω*_*EY*_ = cov(*G*_*E*_,*G*_*Y*_) = 0 using a simulation procedure (see Werme et al. (2022) for details). Note that the LAVA model as described in Werme et al. (2022) is technically defined in terms of testing *H*_0_: cov(*δ*_*E*_,*δ*_*Y*_) = 0, but since cov 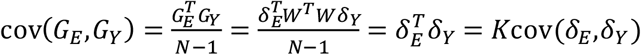 this is equivalent to testing *H*_0_: cov(*G*_*E*_, *G*_*Y*_) = 0.

However, under this null hypothesis 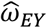 has an expectedvalue of zero and a variance of 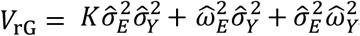. Although it is a sum of product-normal distributions, with large enough *K* its distribution converges on a normal distribution following the central limit theorem, allowing for a normal approximation of the sampling distribution of 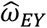. For the simulations and real data analysis we computed LAVA-rG p-values in both ways, and as shown in Supplemental Figure 7 these were found to be virtually identical. As such, all LAVA-rG results reported in this paper are based on p-values computed using the distribution 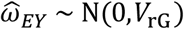, to provide a mathematically more direct comparison for the LAVA-TWAS implementation.

### LAVA implementation of TWAS

To construct a TWAS model within the LAVA framework, we note that in a linear regression for equation (2) we have estimator 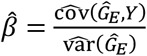. Since 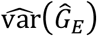 is considered fixed in TWAS the sampling distribution of 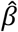 directly proportional to the distribution of 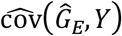, the sample estimate of 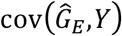. As both 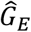 and *Y* have means of zero, and since 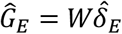, we have 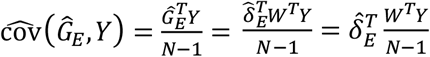. As previously derived 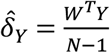, and it therefore follows 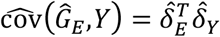. This 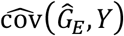 is therefore equal to the genetic covariance estimate 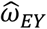 used by LAVA-rG, and thus both LAVA-TWAS and LAVA-rG are shown to use the same test statistic 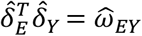.

We can also observe that this 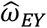 can be rewritten as the sum 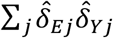, weighting each genetic association 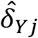 by the corresponding eQTL estimate 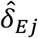. This implementation is therefore analogous to how TWAS is performed using GWAS summary statistics in other TWAS methods (eg. Gusev (2016)^6^), except defined in terms of the estimated genetic associations of principal component matrix *W* rather than the original SNP genotype matrix *X*.

### Data

Genotype data for simulations came from the UK Biobank ^28^, a national genomic study of over 500,000 volunteer participants in the UK. The study was approved by the National Research Ethics Service Committee North West–Haydock (reference 11/NW/0382) and data were accessed under application #16406. Data collection, primary quality control, and imputation of the genotype data were performed by the UK Biobank^29^. Data was converted to hardcalled genotypes, and then additionally filtered, using only SNPs with an info score of at least 0.9 and missingness no greater than 5%, and restricting the sample to unrelated, European-ancestry individuals with concordant sex. Remaining missing genotype values were then mean-imputed to ensure consistency of the input data across different methods.

The European panel of the 1,000 Genomes^43^ data (N = 503, as downloaded from https://ctg.cncr.nl/software/magma) was used as genotype reference data to estimate LD for the real data analysis for both LAVA and FUSION, removing SNP with a minor allele frequency (MAF) below 0.5%. For eQTL data we used the GTEx^34^ data (v8, European subset), for whole blood gene expression. The published cis-eQTL summary statistics (European ancestry) from GTEx Portal were used for the LAVA analysis, pre-computed FUSION model files were used for the FUSION analysis. For every analysed gene, we included all SNPs in the data within one megabase of the transcription start site. Genes were filtered to include only autosomal protein-coding and RNA genes expressed in blood, and only genes also available in the FUSION model files were used, resulting in a total of 14,584 genes available for analysis.

GWAS summary statistics were selected for five well-powered phenotypes, chosen to reflect a range of different domains. These were BMI (GIANT)^30^ (no waist-hip ratio adjustment), educational attainment (SSGAC)^31^, schizophrenia (PGC, wave 3)^32^, diastolic blood pressure (GWAS Atlas)^33^ and type 2 diabetes (GWAS Atlas)^33^. Sample size and number of SNPs for each sample can be found in Table 2.

### Simulation study

Simulation studies were performed to evaluate the type 1 error rates of TWAS when used to evaluate the null hypothesis *H*_0_:cov(*G*_*E*_,*G*_*Y*_) = 0. The simulations were based on the UK Biobank data, generating gene expression and outcome phenotypes while varying their sample sizes and the local heritabilities, as well as the number of SNPs per gene. The number of SNPs *K* in the genes was varied across values of 100, 500, 1,000 and 2,500 SNPs; sample sizes of either 250 or 1,000 were used for the gene expression (denoted *N*_*E*_), and 10,000, 50,000 or 100,000 for the outcome phenotype (denoted *N*_*Y*_); and local heritabilities, defined as the percentage of variance explained by the SNPs in the gene, were set at either 1%, 5% or 10% for the gene expression (denoted 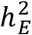)^44^, and 0.001%, 0.005%, 0.01%, 0.05%, 0.1%, 0.5% or 1% for the outcome phenotype (denoted 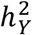). The number of SNPs *K* was only varied at sample size settings of *N*_*E*_ = 1,000 and *N*_*Y*_ = 10,000, and the sample sizes were only varied at *K* = 1,000. All combinations of the two local heritabilities were simulated for each setting of *K* and sample sizes. The simulations were analysed using LAVA-TWAS, LAVA-rG, the four FUSION models and CoMM.

To create the samples to use, we first drew a random subset of 100,000 individuals from the UK Biobank data. For each sample size *N* used in the simulations, the first *N* individuals from this subset were used for that sample, to ensure that samples of different sizes are all nested. The MAF for all SNPs was computed based on the *N* = 100 subsample, discarding all SNPs with MAF below 5%. Since the subsamples are nested, this ensures that all SNPs have a sufficient minor allele count at every sample size used. Ten blocks of 2,500 consecutive SNPs were then selected to represent simulated genes, one block each from around the middle of the first ten chromosomes (avoiding the centromere region). The number of SNPs *K* was varied across conditions by using only a subset of these 2,500 SNPs, always selecting the first *K* SNPs from each block. For each simulation condition, the total number of iterations for that condition was evenly distributed over these ten blocks, and results were aggregated over the blocks. Type 1 error rates per condition were computed as the proportion of iterations for that condition for which the p-value was smaller than the significance threshold, which was set to 0.05 unless otherwise specified.

To simulate gene expression *E* and outcome phenotype *Y* for the standardized genotypes *X* of the SNPs in a gene, *X* was projected onto standardized principal components *W*, pruning away redundant components based on the cumulative genotypic variance explained by the components (retaining those that jointly explain 99% of the total variance). For each iteration, true genetic effect sizes *δ*_*E*_ and *δ*_*Y*_ for the principal components under the null hypothesis of cov(*G*_*E*_,*G*_*Y*_) = 0 were generated by drawing values from a normal distribution, then regressing one vector on the other and retaining only the residuals for the outcome vector to ensure that *δ*_*E*_ and *δ*_*Y*_ were exactly independent. The two genetic components were then computed as *G*_*E*_ = *Wδ*_*E*_ and *G*_*Y*_ = *Wδ*_*Y*_, and both were subsequently standardized to a variance of one.

Simulated gene expression and phenotypevalues were then generated as *E* = *Wδ*_*E*_ + *ξ*_*E*_ and *Y* = *Wδ*_*Y*_ + *ξ*_*Y*_, drawing the residuals *ξ*_*E*_ and *ξ*_*Y*_ from normal distributions with variance parameters equal to 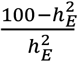 and 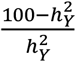 respectively, ensuring the desired level of explained variance. Both *E* and *Y* were then regressed on each individual SNP in *X* using simple linear regression, and the resulting test statistics as well as the simulated *E* and *Y* themselves were stored for subsequent analysis.

To compare the different methods, a baseline set of simulation conditions with 1,000 iterations, with *K* = 1,000, *N*_*E*_ = 1,000 and the remaining parameters across their full range, was generated and analysed with all methods: LAVA-TWAS, LAVA-rG, FUSION-BLUP, FUSION-BSLMM, FUSION-ElastNet, FUSION-LASSO and CoMM. LAVA analyses were performed using the summary statistics as input, FUSION analyses with the raw gene expression values for the first stage and outcome phenotype summarystatistics for input, and CoMM using the raw values for both gene expression and outcome phenotype. For the first stage of FUSION, cross-validation was turned off and the true 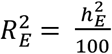 value was set using the --hsq_set option. In the CoMM analyses, the model was found to run into convergence issues due to *K* being equal to *N*_*E*_, and the simulations were rerun setting *K* to 100 instead to resolve this. Moreover, an additional set of simulations was performed with 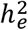 values of 1%, 2%, 4%, 5%, 6%, 8% and 10%, and 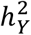 values of 0.01%, 0.02%, 0.05%, 0.1%, 0.2%, 0.5% and 1%.

For LAVA-TWAS, further of simulations with 10,000 iterations were performed across the full range of parameter settings for analysis, to evaluate the impact of these parameters on type 1 error rates. To determine the effect of filtering genes on the presence of eQTL signal, as is common practice in TWAS analyses, for these simulations we also computed the LAVA univariate p-value for the gene expression. In addition to the regular type 1 error rates, filtered type 1 error rates were also computed based on only the iterations for which the univariate p-value was below either 0.05 or 0.0001. Finally, to allow for reliable computation of type 1 error rates at lower significance thresholds, for the subset of these conditions with *K* = 1,000, *N*_*E*_ = 1,000, 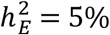 and 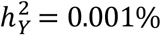, 0.01%, 0.1% or 1% the number of iterations was increased to 100,000. Note that although for LAVA-TWAS the number of iterations differs from the other methods and varies across conditions, for each plot in this paper the type 1 error rate values within that plot are always based on the same number of iterations.

Additional simulations were also performed under the TWAS null hypothesis of 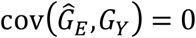 to validate the LAVA-TWAS model. To do so, the above procedure was modified to first generate *δ*_*E*_ and the corresponding *G*_*E*_, from which *E* was then simulated. This was used to estimate SNP summary statistics for the associations between *X* and *E*, from which the estimate vector 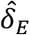 was then computed for the outcome data. Finally, *δ*_*Y*_was then generated to be exactly independent of this 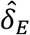, with the rest of the simulation process proceeding as normal. For 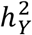, the parameter range was extended to also include 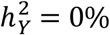.

### Real data analysis

TWAS and local genetic correlation analyses were performed on the GWAS data as follows, separately for each of the five phenotypes. In all the analyses, SNP filtering was applied to remove all SNPs with a minor allele frequency below 0.5%, and only SNPs available in the 1,000 Genomes data, the GTEx data, and the GWAS sample for that phenotype were used.

For every gene, univariate LAVA analysis for the eQTL signal, LAVA-rG and LAVA-TWAS analyses, and FUSION analyses with the Elastic Net and LASSO models was performed. For FUSION,the BLUP and BSLMM models were not used as these were not available in the precomputed model files. FUSION univariate p-values for the eQTL signal were also obtained from these model files. Note that separate SNP filtering was applied prior to these models being computed, and as such the LAVA and FUSION analyses are based on different sets of SNPs per gene.

For the primary analyses, the bivariate LAVA and FUSION analyses were only performed for genes for which the LAVA univariate tests were significant after correcting for the total number of genes, at a threshold of 0.05/14,584 = 3.43 × 10^−6^ (see Supplemental Table 1 for results when using the FUSION univariate p-values instead). The significance threshold for the bivariate analyses was set separately for each phenotype, at a Bonferroni-correction for the total number of genes that was univariate significant for that phenotype (see Table 2). In a secondary analysis, to evaluate TWAS results in the absence of gene expression signal, bivariate analyses were performed for all genes for which the LAVA univariate p-value was greater than 0.05, again setting the significance threshold per phenotype for the number of genes with univariate p-value greater than 0.05 for that phenotype.

## Data availability

For the real data analysis, summary statistics were obtained for five different phenotypes: blood pressure (https://atlas.ctglab.nl/traitDB/3379), BMI (https://zenodo.org/record/1251813#.YBAfYBbvJhE, combined.23May2018), educational attainment (https://www.thessgac.org/data), schizophrenia (https://www.med.unc.edu/pgc/download-results/, scz2022), type 2 diabetes (https://atlas.ctglab.nl/traitDB/3328). GTEx v8 European ancestry whole blood eQTL summary statistics and the gene definition file used for the LAVA-TWAS and LAVA-rG analyses were obtained via https://www.gtexportal.org/home/datasets, the corresponding FUSION model files from http://gusevlab.org/projects/fusion/#single-tissue-gene-expression (EUR Samples). The European subset of the 1000 Genomes data used as LD reference for the real data analyses was downloaded from https://ctg.cncr.nl/software/magma. UK Biobank genotype data was obtained from https://www.ukbiobank.ac.uk/ under application #16406.

## Code availability

LAVA is implemented as an R package, which is publicly available at the LAVA website (https://ctg.cncr.nl/software/lava) and LAVA GitHub repository (https://github.com/josefin-werme/LAVA). GTEx v8 input files compatible with LAVA are available from the LAVA website as well. Version 0.1.0 of LAVA was used for the LAVA analyses in this paper; this version contains specific functionality for handling eQTL summary statistics input files. FUSION can be obtained from http://gusevlab.org/projects/fusion/ and CoMM from https://github.com/gordonliu810822/CoMM. Scripts used for the simulations and analyses performed for this paper are available from https://github.com/cadeleeuw/twas-validity-scripts2022.

## Acknowledgements

This work was funded by The Netherlands Organization for Scientific Research (Grant No. NWO VICI 435–14–005 [to DP]) and NWO Gravitation: BRAINSCAPES: A Roadmap from Neurogenetics to Neurobiology (Grant No. 024.004.012 [to DP]), and a European Research Council advanced grant (Grant No, ERC-2018-AdG GWAS2FUNC 834057 [to DP, and funding CdL]). CdL was funded by F. Hoffmann-La Roche AG. WJP was funded by an NWO Veni grant (NWO: 916-19-152). JES was supported by an NWO Veni grant (NWO: 201G-064). The research has been conducted using the UK Biobank Resource (application no. 16406). The analyses were carried out on the Genetic Cluster Computer, which is financed by the Netherlands Organization for Scientific Research (NWO: 480-05-003), by the VU University (Amsterdam, The Netherlands) and the Dutch Brain Foundation, hosted by the Dutch National Computing and Networking Services SurfSARA.

## Author contributions

CdL and JW conceived of the study. CdL performed the analyses and simulations. CdL wrote the manuscript, in collaboration with JW. All authors participated in the interpretation of the results and revision of the manuscript, and provided meaningful contributions at each stage of the project.

## Competing interests

The authors declare no competing financial interests.

## Notes

### Competing Interest Statement

The authors have declared no competing interest.

### Summary of Updates

Revised the main text to clarify the core points, and added some additional simulation results.

