## Supplemental Methods for "On the interpretation of transcriptome-wide association studies"

### Table of Contents

**The TWAS null value** 2

**Underlying distributions** 2

**Non-linear TWAS models** 4

**Comparison with the CoMM model** 5

**FUSION permutation test** 5

**Discussion of simulation results** 6

**Supplemental figures** 8

*Supplemental Figure 1. Type 1 error rate simulation results as a function of the number of SNPs in the gene* 8

*Supplemental Figure 2. Type 1 error rate simulation results as a function of univariate input filtering* 9

*Supplemental Figure 3. Type 1 error rate simulation results for LAVA-TWAS under the TWAS null hypothesis* 10

*Supplemental Figure 4. Type 1 error rate simulation results for FUSION models* 11

*Supplemental Figure 5. Type 1 error rate simulation results for CoMM.* 12

*Supplemental Figure 6. P-value bias in the real data analysis for BMI.* 13

*Supplemental Figure 7. Comparison of -log10 p-values for different implementations of LAVA-rG.* 13

*Supplemental Figure 8. Type 1 error rate simulation results for FUSION models with post hoc permutation test* 14

**Supplemental tables** 15

*Supplemental Table 1. Summary of results of TWAS and local genetic correlation analyses of published summary statistics for five phenotypes, with FUSION univariate filtering.* 15

*Supplemental Table 2. Summary of results of TWAS and local genetic correlation analyses for genes with no detectable eQTL signal.* 15

*Supplemental Table 3. Summary of results of FUSION permutation tests for published summary statistics for five phenotypes.* 15

**Supplemental references** 16

##### THE TWAS NULL VALUE

As shown in the main text, TWAS can be interpreted as performing a test of the local genetic covariance $H_0\colon \mathrm{cov}(G_E, G_Y) = -\hat{C}$, raising the question of how this value $\hat{C}$ can be interpreted. The $\hat{C} = \mathrm{cov}(\hat{\Delta}_E, G_Y)$ term arises in the decomposition $\mathrm{cov}(\hat{G}_E, G_Y) = \mathrm{cov}(G_E + \hat{\Delta}_E, G_Y) = \mathrm{cov}(G_E, G_Y) + \mathrm{cov}(\hat{\Delta}_E, G_Y) = \mathrm{cov}(G_E, G_Y) + \hat{C}$, showing that in a general sense it can be seen as an error term in $\mathrm{cov}(\hat{G}_E, G_Y)$ if this were used as an estimator of the true local genetic covariance $\mathrm{cov}(G_E, G_Y)$. This itself in turn arises from the decomposition of $\hat{G}_E$ as $\hat{G}_E = G_E + \hat{\Delta}_E$, with $\hat{\Delta}_E$ again an error term relative to the true genetic component $G_E$. Finally, if we similarly decompose the estimated genetic effects on the gene expression $\hat{\alpha}_E$ into $\hat{\alpha}_E = \alpha_E + \hat{w}$, with $\hat{w}$ the error in the estimation of $\alpha_E$, we can see that $\hat{G}_E = X\hat{\alpha}_E = X(\alpha_E + \hat{w}) = X\alpha_E + X\hat{w} = G_E + X\hat{w}$, and therefore $\hat{\Delta}_E = X\hat{w} = \sum_j X_j \hat{w}_j$.

Plugging this back into $\hat{C} = \mathrm{cov}(\hat{\Delta}_E, G_Y)$, we find that $\hat{C} = \mathrm{cov}(\sum_j X_j \hat{w}_j, G_Y) = \sum_j \hat{w}_j \mathrm{cov}(X_j, G_Y) = \sum_j \hat{w}_j \mathrm{cov}(X_j, Y)$, where the last step follows from the fact that $\mathrm{cov}(X_j, Y) = \mathrm{cov}(X_j, G_Y + \xi_Y) = \mathrm{cov}(X_j, G_Y) + \mathrm{cov}(X_j, \xi_Y) = \mathrm{cov}(X_j, G_Y)$ for every SNP $j$, since by definition the $\xi_Y$ term from equation (3) is independent of $X$. The exact process by which the values of $\hat{w}$ are set will depend on the methods used to estimate equation (1). For example, in the simple case of Ordinary Least Squares multiple linear regression, $\hat{w}$ is a draw from a $\mathrm{MVN}(0, \eta_E^2 (X^T X)^{-1})$, with $\eta_E^2 = \mathrm{var}(\xi_E)$. For other methods such as elastic net or LASSO, the distribution from which $\hat{w}$ is drawn will be more complex, and may not have a closed-form expression. Nevertheless, regardless of the exact form of this distribution, $\hat{w}$ will still be the product of such a random process resulting from the fact that $\alpha_E$ is estimated from a finite sample, and the value of $\hat{w}$ will therefore be largely or wholly independent of $\mathrm{cov}(X, Y)$.

Because there is no systematic relationship between $\hat{w}_j$ and $\mathrm{cov}(X_j, Y)$, and the values of $\hat{w}_j$ are randomly drawn and sample-specific, rather than population-level parameters, the value of $\hat{C} = \sum_j \hat{w}_j \mathrm{cov}(X_j, Y)$ is itself random and sample-specific, its value fixed when the gene expression sample was generated. If there is any genetic between the SNPs in $X$ and the phenotype $Y$, for $\hat{C}$ to equal $-\mathrm{cov}(G_E, G_Y)$ the weights $\hat{w}_j$ would have to be such that by chance the sum $\sum_j \hat{w}_j \mathrm{cov}(X_j, Y)$ equals $-\mathrm{cov}(G_E, G_Y)$, the probability of which is negligible in practice. Realistically, the only scenario in which $\hat{C}$ will equal $-\mathrm{cov}(G_E, G_Y)$ and the TWAS null hypothesis will be true, is if there is if $X$ is independent of $Y$, ie. $\mathrm{cov}(X, Y) = 0$. In this case, $\hat{C}$ will be zero regardless of the value of $\hat{w}$. Moreover, by virtue of the fact that absent any genetic signal the variance of $G_Y$ will then also be zero, and consequently the corresponding covariance $\mathrm{cov}(G_E, G_Y)$ must be zero as well.

##### UNDERLYING DISTRIBUTIONS

The fundamental cause of the issue with TWAS is that the data generation encompasses two separate sampling processes, but the model in equation (2) that is tested only accounts for one, the sampling of the phenotype. The sampling of the gene expression is omitted, conditioning on $\hat{G}_E$ rather than modeling its full distribution around $G_E$. In this section we will outline the distinction between these joint and conditional distributions, and how this relates to the issue described. We will do so from the perspective of the LAVA-TWAS model, as it can be directly compared to LAVA-rG, and as the resulting distributions are mathematically much more tractable than those of other stage 1 fitting approaches like the LASSO or Elastic Net implemented in FUSION. Although the specific distributions will therefore be particular to LAVA-TWAS, the same principles apply to any other traditional TWAS method that estimates and then conditions on $\hat{G}_E$.

In the LAVA framework, we first define the standardized principal component matrix $W$ of $X$, such that $G_E = X\alpha_E = W\delta_E$ and $G_Y = X\alpha_Y = W\delta_Y$, which simplifies the subsequent distributions by dealing with the LD in the projection from $X$ to $W$, without loss of information. Multiple linear regression for $E$ then yields a sampling distribution $\hat{\delta}_E \sim \mathrm{MVN}(\delta_E, \sigma_E^2 I_K)$ with $\sigma_E^2 = \frac{\eta_E^2}{N_E - 1}$ the sampling variance $\eta_E^2$ the residual variance, $N_E$ the sample size, and $K$ the number of principal components, and likewise $\hat{\delta}_Y \sim \mathrm{MVN}(\delta_Y, \sigma_Y^2 I_K)$. Assuming no overlap between the samples, these two distributions are independent. Local genetic variances are $\omega_E^2 = \mathrm{var}(G_E) = \delta_E^T \delta_E$ and $\omega_Y^2 = \mathrm{var}(G_Y) = \delta_Y^T \delta_Y$, with local genetic covariance $\omega_{EY} = \mathrm{cov}(G_E, G_Y) = \delta_E^T \delta_Y$. The variances can be estimated using the method of moments as $\hat{\omega}_E^2 = \hat{\delta}_E^T \hat{\delta}_E - K\hat{\sigma}_E^2$ and $\hat{\omega}_Y^2 = \hat{\delta}_Y^T \hat{\delta}_Y - K\hat{\sigma}_Y^2$, and the covariance as $\hat{\omega}_{EY} = \hat{\delta}_E^T \hat{\delta}_Y = \sum_j \hat{\delta}_{Ej} \hat{\delta}_{Yj}$ (see also *Methods – Local genetic correlation*), which is also used as the test statistic for both LAVA-rG and LAVA-TWAS to test their respective null hypotheses.

Under the generative model, the individual elements $\hat{\delta}_{Ej}\hat{\delta}_{Yj}$ have an expected value $E[\hat{\delta}_{Ej}\hat{\delta}_{Yj}] = E[\hat{\delta}_{Ej}]E[\hat{\delta}_{Yj}] = \delta_{Ej}\delta_{Yj}$ and a variance $\mathrm{var}(\hat{\delta}_{Ej}\hat{\delta}_{Yj}) = \delta_{Ej}^2\sigma_Y^2 + \delta_{Yj}^2\sigma_E^2 + \sigma_E^2\sigma_Y^2$. Consequently, $E[\hat{\delta}_E^T\hat{\delta}_Y] = \sum_j E[\hat{\delta}_{Ej}\hat{\delta}_{Yj}] = \delta_E^T\delta_Y = \omega_{EY}$, and $\mathrm{var}(\hat{\delta}_E^T\hat{\delta}_Y) = \sum_j \mathrm{var}(\hat{\delta}_{Ej}\hat{\delta}_{Yj}) = \delta_E^T\delta_E\sigma_Y^2 + \delta_Y^T\delta_Y\sigma_E^2 + K\sigma_E^2\sigma_Y^2 = \omega_E^2\sigma_Y^2 + \omega_Y^2\sigma_E^2 + K\sigma_E^2\sigma_Y^2$. Given sufficiently large $K$, which will generally apply with the typical number of SNPs per gene in a TWAS setting, following the central limit theorem the distribution of $\hat{\delta}_E^T\hat{\delta}_Y$ can be closely approximated with a normal distribution. For simplicity, we will therefore assume that $\hat{\delta}_E^T\hat{\delta}_Y$ is normally distributed in the remainder of this section.

Plugging in the sample estimates of the parameters in the variance, this thus yields a marginal distribution $\hat{\delta}_E^T\hat{\delta}_Y \sim \mathrm{N}(\omega_{EY}, \hat{\omega}_E^2\hat{\sigma}_Y^2 + \hat{\omega}_Y^2\hat{\sigma}_E^2 + K\hat{\sigma}_E^2\hat{\sigma}_Y^2)$ with which the null hypothesis $H_0\colon \mathrm{cov}(G_E, G_Y) = \omega_{EY} = 0$ can be tested, as in LAVA-rG. By contrast however, TWAS works with the conditional distribution of its test statistic given $\hat{G}_E$, so in the case of LAVA-TWAS this would be the distribution of $\hat{\delta}_E^T\hat{\delta}_Y$ given $\hat{\delta}_E$. This is simply a linear transformation of the multivariate normal distribution of $\hat{\delta}_Y$, and as such has an expected value $E[\hat{\delta}_E^T\hat{\delta}_Y|\hat{\delta}_E] = \hat{\delta}_E^T\delta_Y = \mathrm{cov}(\hat{G}_E, G_Y) = \mathrm{cov}(G_E, G_Y) + \hat{C}$ and a variance of $\mathrm{var}(\hat{\delta}_E^T\hat{\delta}_Y|\hat{\delta}_E) = \hat{\delta}_E^T\hat{\delta}_E\hat{\sigma}_Y^2 = \hat{\omega}_E^2\hat{\sigma}_Y^2 + K\hat{\sigma}_E^2\hat{\sigma}_Y^2$, where the second step follows from the fact that $\hat{\delta}_E^T\hat{\delta}_E = \hat{\omega}_E^2 + K\hat{\sigma}_E^2$. Thus, the conditional sampling distribution used by LAVA-TWAS is $\hat{\delta}_E^T\hat{\delta}_Y|\hat{\delta}_E \sim \mathrm{N}(\hat{\delta}_E^T\delta_Y, \hat{\omega}_E^2\hat{\sigma}_Y^2 + K\hat{\sigma}_E^2\hat{\sigma}_Y^2)$, which under its null hypothesis $H_0\colon \mathrm{cov}(\hat{G}_E, G_Y) = \hat{\delta}_E^T\delta_Y = 0$ therefore reduced to $\hat{\delta}_E^T\hat{\delta}_Y|\hat{\delta}_E \sim \mathrm{N}(0, \hat{\omega}_E^2\hat{\sigma}_Y^2 + K\hat{\sigma}_E^2\hat{\sigma}_Y^2)$.

From this we can see that LAVA-rG and LAVA-TWAS evaluate their respective null hypotheses by comparing the same test statistic $\hat{\delta}_E^T\hat{\delta}_Y$ against a normal distribution with a mean of zero and with variances of $V_{\mathrm{rG}} = \hat{\omega}_E^2\hat{\sigma}_Y^2 + \hat{\omega}_Y^2\hat{\sigma}_E^2 + K\hat{\sigma}_E^2\hat{\sigma}_Y^2$ and $V_{\mathrm{TWAS}} = \hat{\omega}_E^2\hat{\sigma}_Y^2 + K\hat{\sigma}_E^2\hat{\sigma}_Y^2$. Since $V_{\mathrm{rG}} = V_{\mathrm{TWAS}} + \hat{\omega}_Y^2\hat{\sigma}_E^2$, inherently $V_{\mathrm{rG}}$ will be larger than $V_{\mathrm{TWAS}}$; and thus so will the standard errors, as these are simply the square roots of $V_{\mathrm{rG}}$ and $V_{\mathrm{TWAS}}$ respectively. Specifically, we can express the deflation of the standard errors as $\sqrt{V_{\mathrm{TWAS}}/V_{\mathrm{rG}}} = \sqrt{1 - \hat{\omega}_Y^2\hat{\sigma}_E^2/(\hat{\omega}_E^2\hat{\sigma}_Y^2 + \hat{\omega}_Y^2\hat{\sigma}_E^2 + K\hat{\sigma}_E^2\hat{\sigma}_Y^2)}$.

If our aim is to evaluate $H_0\colon \mathrm{cov}(G_E, G_Y) = 0$, testing for the presence of a local genetic relationship, then given the generative model described above we must use the marginal distribution of $\hat{\delta}_E^T\hat{\delta}_Y$ rather than its the conditional distribution given $\hat{\delta}_E$, since we know that the gene expression $E$ was itself sampled, and this sampling process induces additional uncertainty in our estimates that must be accounted for. Given the choice of $\hat{\delta}_E^T\hat{\delta}_Y$ as test statistic, this can therefore only be achieved by using the sampling distribution $\hat{\delta}_E^T\hat{\delta}_Y \sim \mathrm{N}(0, V_{\mathrm{rG}})$, since this is the distribution directly implied by the full generative model.

As such, if we were to incorrectly interpret and use LAVA-TWAS as if it tests $H_0\colon \mathrm{cov}(G_E, G_Y) = 0$, and thus obtain our p-value by comparing $\hat{\delta}_E^T\hat{\delta}_Y$ to the TWAS sampling distribution $\mathrm{N}(0, V_{\mathrm{TWAS}})$ instead, this p-value will be too, low since $V_{\mathrm{TWAS}} < V_{\mathrm{rG}}$. Because the only difference between LAVA-rG and LAVA-TWAS is the shift from using the full generative model to conditioning on the uncertainty due to the sampling of $E$, with both the test statistic used and the generative model otherwise completely identical, any results declared significant when using $\mathrm{N}(0, V_{\mathrm{TWAS}})$ that wouldn't be significant when using $\mathrm{N}(0, V_{\mathrm{rG}})$, must inherently be spurious. In cases where $H_0\colon \mathrm{cov}(G_E, G_Y) = 0$ is true, this will result in an inflation of the type 1 error rate, whereas in cases where $H_0\colon \mathrm{cov}(G_E, G_Y) = 0$ it will lead to results being invalid because the statistical evidence in the data is not sufficient to support that conclusion.

For other methods of fitting equation (1) and obtaining $\hat{G}_E$ the distributions involved will be different, but the same principles still apply: the estimates from which $\hat{G}_E$ is computed are obtained from a finite sample and are therefore subject to uncertainty, and in the traditional TWAS methods this uncertainty is conditioned on in equation (2), rather than having its full distribution accounted for in the model. And because a source of variance that is known to exist in the data generation process is omitted, when interpreting the TWAS results as if they were a test of $H_0\colon \mathrm{cov}(G_E, G_Y) = 0$, the standard errors will be deflated.

**NON-LINEAR TWAS MODELS**

Several of the methods listed in Table 1 ('non-linear models') more explicitly approach TWAS as a form of joint association testing, and seek to generalize the model to relax this expectation of consistent effect directions, as follows. For a linear regression based on equation (2), the corresponding test statistic can be expressed as $T_1 = c_1 \sum_j w_j r_j$, with $r_j$ the correlation between SNP $j$ and the phenotype, and $w_j$ the corresponding (signed) weight derived from the eQTL data (and $c_1$ a scaling constant). This follows from the fact that $\mathrm{cov}(\hat{G}_E, Y) = \sum_j \hat{\alpha}_{Ej} \mathrm{cov}(X_j, Y)$).

If the signs of $w_j$ and $r_j$ are either all the same or all opposite, each term $w_j r_j$ will have the same sign, and the test statistic as a whole will tend to have a large value. If the signs are not consistent however, the $w_j r_j$ for different SNPs will have different signs, and they (partially) cancel each other out when summed. To get around this, we can define a generalized test statistic $T_P = c_P \sum_j (w_j r_j)^P$, raising the terms $w_j r_j$ to the power $P$ before summing them. This has the dual effect of giving more weight in the test statistic to SNPs with stronger correlations with $Y$, as well as removing the expectation of consistent effect directions for even values of $P$. A notable special case of this test statistic is $T_2$, which yields the test statistic of the weighted SKAT model[1].

The different methods in the 'non-linear' section of Table 1 use this test statistic in different ways (with all the 'linear' methods using only $T_1$). Tang (2021)[2] uses $T_2$ instead of $T_1$, Zhang (2020)[3] uses an adaptive combination of $T_1$ and $T_2$, and Xu (2017)[4] uses an adaptive combination of $T_1, T_2, \ldots, T_6, T_\infty$ (where $T_\infty = \max(|w_j r_j|)$. Since all three of these methods include $T_2$, they have better power to detect joint associations when effect sizes are not consistent with $\hat{\alpha}_E$, though at the expense of somewhat lower power in scenarios where the linear model is a good fit. However, these tests can no longer be defined in terms of $\mathrm{cov}(\hat{G}_E, G_Y)$, much less $\mathrm{cov}(G_E, G_Y)$, making the results even more difficult to interpret even if the main issue discussed in this paper were addressed.

Moreover, the null hypothesis for all three models implies that no genetic associations exist between the phenotype and the SNPs included in the analysis. Tang (2021) is defined using the linear equation $Y = X\alpha_Y + \xi_Y$ in (3), like the traditional linear models, integrating eQTL-based weights by defining $\alpha_Y \sim \mathrm{MVN}(0, \tau^2 W)$, where $W$ is a diagonal matrix with the squared weights on the diagonal. It tests the null hypothesis $H_0\colon \tau^2 = 0$, reducing the null model to $Y = \xi_Y$, implying $Y$ is independent of $X$ under the null. For the other two methods, null assumptions are placed directly on the vector of score statistics $X^T Y$ (Xu, 2017) or the corresponding SNP Z-statistics (Zhang, 2020), drawing these from a centered multivariate normal distribution (with covariance matrix a function of the LD matrix and standard errors for the SNP genetic associations) that similarly implies absence of genetic association of these SNPs with the phenotype. Like the traditional linear TWAS models, these non-linear models thus operate as essentially a form of joint genetic association analysis for the phenotype.

#### Comparison with the CoMM model

Unlike all the other TWAS methods in Table 1, CoMM[5] explicitly accounts for the uncertainty in the eQTL estimates. It accomplishes this by specifying the linear equations (translating to our notation) $E = X\alpha + \xi_E$ and $Y = X\alpha\beta + \xi_Y$ (with the two equations sharing the same parameter $\alpha$), which, if we substitute $G = X\alpha$, translates to $E = G + \xi_E$ and $Y = G\beta + \xi_Y$, analogous to equations (1) and (2) for the regular TWAS framework. Note however that as in equation (3), the residual term $\xi_Y$ is assumed to be independent of $X$ in general, in contrast to the residual $\varepsilon_Y$ in (2) which is only assumed to be independent of $X\hat{\alpha}_E$. The parameters $\alpha$ and $\beta$ are estimated simultaneously in a single model using both these equations, and by not using a separately estimated $\hat{G}$ it avoids the problem in the other TWAS methods of failing to account for the estimation uncertainty in that $\hat{G}$. A test of $H_0: \beta = 0$ is performed for each gene to obtain an output p-value.

Although CoMM seems to address the issue with TWAS discussed in the main text, the problem with this model is that it uses the same parameter $\alpha$ to capture the genetic associations of $X$ with both $E$ and $Y$, in contrast to equations (1) and (3) which provide separate parameter vectors $\alpha_E$ and $\alpha_Y$. In effect, the CoMM model is equivalent to simultaneously fitting (1) and (2), with $\alpha_E = \alpha$ and $\alpha_Y = \alpha\beta = \alpha_Y\beta$. As such, it essentially imposing a very strong constraint on the genetic effects of the SNPs in $X$, assuming that the effects of $X$ on $E$ are proportional to the effects of $X$ on $Y$.

The issue with this becomes readily apparent when we inspect the model under the null hypothesis $H_0: \beta = 0$. Under that null, the first equation $E = X\alpha + \xi_E$ is unchanged since $\beta$ does not occur in it, but for $Y$ we obtain $Y = X\alpha\beta + \xi_Y = X\alpha \times 0 + \xi_Y = \xi_Y$. As noted, this residual term is entirely independent of $X$, and this null model therefore implies that there are no genetic associations between $X$ and $Y$ at all. In other words, this null hypothesis is again equivalent to testing $\alpha_Y = 0$, reducing to a joint association test just as the other TWAS models do.

#### FUSION permutation test

FUSION[6] implements an additional, post-hoc permutation test that can optionally be applied when the primary TWAS test is significant, which works by randomly permuting $\hat{\alpha}_E$ to obtain an empirical p-value. The exact null hypothesis evaluated by this test is left implicit, and the fact that the permutations ignore LD make it difficult to relate it directly to the local genetic covariance $\mathrm{cov}(G_E, G_Y)$, but nevertheless we aimed to evaluate in both our simulations and real data analysis whether it might help address the issue with TWAS raised in this paper.

When running this test, p-values were based on at least 1,000 permutations. For the real data analysis, the number of permutations was increased using an adaptive procedure, modifying the procedure built into FUSION. The number of permutations was increased ten-fold each step, until either the squared Z-statistic for at least 10 permutations was higher than the observed squared Z-statistic, or a maximum of 10,000,000 was reached.

Results in our simulations showed that except for the BLUP model, type 1 error rates under the $H_0: \mathrm{cov}(G_E, G_Y) = 0$ null hypothesis were generally well-controlled at a nominal significance threshold of 0.05, especially when using the permutation test in conjunction with the primary TWAS test, rather than on its own (Supplemental Figure 8). However, when using the permutation test in the real data, at multiple testing corrected significance threshold, it showed a severe reduction in the number of significant genes, with few if any remaining (Supplemental Table 3). As these numbers are also far lower than those found by LAVA-rG, this suggests that the permutation test is poorly powered to detect genetic relationships between gene expression and the phenotype, likely related to the fact that the permutation procedure does not account for the LD between SNPs. Although the FUSION permutation test thus does seem to control type 1 error rates relative to the $H_0: \mathrm{cov}(G_E, G_Y) = 0$ null hypothesis of no local genetic relationship, the severely depleted power means that it nevertheless lacks practical utility.

### DISCUSSION OF SIMULATION RESULTS

The main simulation results for LAVA-TWAS are depicted in Figure 1, and one of the most striking features is the strong effect of both sample size and local heritability for the outcome phenotype. One way to understand this is in light of the fact that, as discussed, TWAS effectively performs a form of joint association test. With greater local heritability for $Y$ there is more effect in the $\alpha_Y$ vector to detect, and with a larger sample size the test has more power to detect them. Under the null hypothesis of $H_0\colon \mathrm{cov}(G_E, G_Y) = 0$ this therefore translates to increasing inflation of the type 1 error rate as a function of these parameters.

Although much less pronounced, Figure 1 also shows an opposite effect on type 1 error rates for the sample size and local heritability of the gene expression. This is due to the fact that with increasing sample size the estimated effect vector $\hat{\alpha}_E$ will converge ever closer on $\alpha_E$, meaning that $\hat{G}_E$ converges on $G_E$ and therefore $\mathrm{cov}(\hat{G}_E, G_Y)$ on $\mathrm{cov}(G_E, G_Y)$. This process is further enhanced by greater local heritability of the gene expression, since this means that the relative amount of noise is decreased since $\xi_E$ in equation (1) now accounts for a smaller proportion of the variance of $E$.

Supplemental Figure 1 shows an interaction between the number of SNPs in the gene and the heritability of the outcome phenotype. This is likely due to two separate competing processes, with on the gene expression side more SNPs in the analysis increasing overfitting and therefore deviation of $\hat{G}_E$ from $G_E$, leading to greater type 1 error rate inflation with more SNPs included; while some other process on the outcome phenotype side is causing type 1 error rate inflation when the number of SNPs is reduced, interacting with the overall level of genetic signal $h_Y^2$ for the outcome. This would account for the fact that, due to the first process, the type 1 error rate starts out higher with more SNPs, but as the $h_Y^2$ goes up the second process starts to become dominant and the trend eventually reverses.

For this latter process probably relates to the fact that as noted, the term $\hat{C}$ that drives the deviation from $H_0\colon \mathrm{cov}(G_E, G_Y) = 0$ equals the sum $\sum_j \widehat{w}_j \mathrm{cov}(X_j, Y)$. This sum becomes greater the better the $\widehat{w}_j$ and $\mathrm{cov}(X_j, Y)$ values align in terms of their relative size and relative direction. This is a chance process however, and with more SNPs it becomes less likely for those values to consistently align as well, while simultaneously the same level of overall effect is distributed over more SNPs, and therefore lower on average per SNP. As a result, the overall sum would tend to be smaller with more SNPs included. It would require further simulation to determine the full nature of this process, however.

As shown in Figure 2, the error rate inflation also gets progressively worse at lower significance thresholds. The reason for this is that the relative difference between the $N(0, V_{\mathrm{rG}})$ and $N(0, V_{\mathrm{TWAS}})$ gets larger the further we move into the tails of those distributions. This is illustrated in the right panel of Figure 4, showing the output p-values of testing using standard errors at different levels of deflation as a function of the correct reference p-value. As shown, the discrepancy between the reference p-value (black) and the p-value that results from using the wrong distribution (shades of red) gets larger as the reference p-value gets smaller, which translates to a correspondingly greater discrepancy in type 1 error rates as well since this is a function of the horizontal distance with the black reference line at an "Output p-value" value equal to the significance threshold.

Shown in Supplemental Figure 2 are simulation results when adding a filtering step on the univariate genetic signal for the gene expression, as is generally done in TWAS analyses, aiming to reduce multiple testing burden by removing genes for which no discernible genetic component is present anyway. As such, these type 1 error rates are computed by first subsetting the iterations to those passing the univariate p-value threshold (0.05, 0.001 or 0.0001), and computing the type 1 error rate as the proportion of iterations in that subset for which the TWAS p-values are below 0.05. As shown, the type 1 error rate becomes further inflated.

This can be explained as follows. Since $\mathrm{cov}(G_E, G_Y) = 0$ in these null simulations, $\hat{C} = \mathrm{cov}(\hat{G}_E, G_Y) = \sum_j \hat{\alpha}_{Ej} \mathrm{cov}(X_j, Y)$, where as noted above $\hat{C}$ is the term that drives the type 1 error rate inflation. The univariate test for the gene expression is a test of $H_0\colon \alpha_E = 0$, and the corresponding p-value is going to be a direct function of the estimate $\hat{\alpha}_E$, and in particular of how far this deviates from

0. As such, selecting on a low univariate p-value entails selecting for genes (or iterations, in the simulation context) for which $\hat{\alpha}_E$ has a larger overall value. This in turn means that $\sum_j \hat{\alpha}_{Ej}\mathrm{cov}(X_j, Y)$ will tend to be bigger as well, since the $\hat{\alpha}_{Ej}$ for each SNP will on average be larger as well. And hence the type 1 error rate becomes further inflated.

A comparison of LAVA-TWAS with the four FUSION models is given in Figure 3, and as shown the results are generally quite similar. In the 10% gene expression heritability condition the type 1 error rates are essentially the same for all five TWAS models, though in the 1% gene expression heritability condition the Elastic Net and LASSO models show distinctly lower type 1 error rates than the rest. However, this is largely due to the fact that due to the weak genetic signal for the gene expression in these conditions, these two models frequently failed to converge. This issue mostly disappears at higher gene expression heritability values; as shown in Supplemental Figure 4, type 1 error rate inflation for the 5% and 10% gene expression heritability conditions are essentially the same, with only the 1% showing lower levels.

Finally, the top row of Supplemental Figure 5 shows the main simulation results for CoMM, with considerably more severe type 1 error rate inflation than the traditional TWAS models. Although both CoMM and the traditional TWAS methods can be seen as performing a joint association test with $H_0\colon \alpha_Y = 0$ on the model in equation (3), whereas the latter impose a hard constraint on $\alpha_Y$ in their model, CoMM only imposes a soft one. Under the traditional models, $\alpha_Y = \hat{\alpha}_E\beta$, with $\hat{\alpha}_E$ estimated and fixed in in the first stage of the analysis. This therefore imposes the assumption in the second stage that $\alpha_Y$ is strictly proportional to the given effect vector $\hat{\alpha}_E$, leaves the model only the single parameter $\beta$ to maximize the model fit.

By contrast, CoMM assumes $\alpha_Y = \alpha_E\beta$ and estimates both $\alpha_E$ and $\beta$ simultaneously. This means that the model can also use the parameter vector $\alpha_E$ to optimize the fit of the relationship between $X$ and $Y$. Indeed, the estimate of the parameter $\alpha_E$ in this model will be a compromise between the fit for equation (1) and that for equation (3), with the relative weight that the data for expression and outcome have in this compromise dependent on their genetic signal strength, ie. the combination of their sample size and local heritability. Hence, as the signal strength for the outcome phenotype get stronger, the model moves progressively closer to behaving like a regular multiple linear regression model applied to just equation (3).

Of additional note is that the effect of the gene expression heritability is partially inverted, with type 1 error rates in the 5% condition higher than both the 1% and 10% conditions, which is confirmed by the additional simulations at more granular heritability values shown in the bottom row of Supplemental Figure 5. The cause of this is not entirely clear, but it is likely related to the fact that CoMM is simultaneously fitting two equations, $E = X\alpha + \xi_E$ and $Y = X\alpha\beta + \xi_Y$. Under the simulations model with $\mathrm{cov}(G_E, G_Y) = 0$ the constraint of $\alpha_Y = \alpha_E\beta$ does not hold, and as a result the fit for these two equations will compete with each other, since better fit for $E$ comes at the expense of better fit for $Y$. Since what is maximized is a weighted combination of the fit for each, the resulting dynamic is likely responsible for the observed reverse in the effect of the heritability of the gene expression on the type 1 error rate.

### SUPPLEMENTAL FIGURES

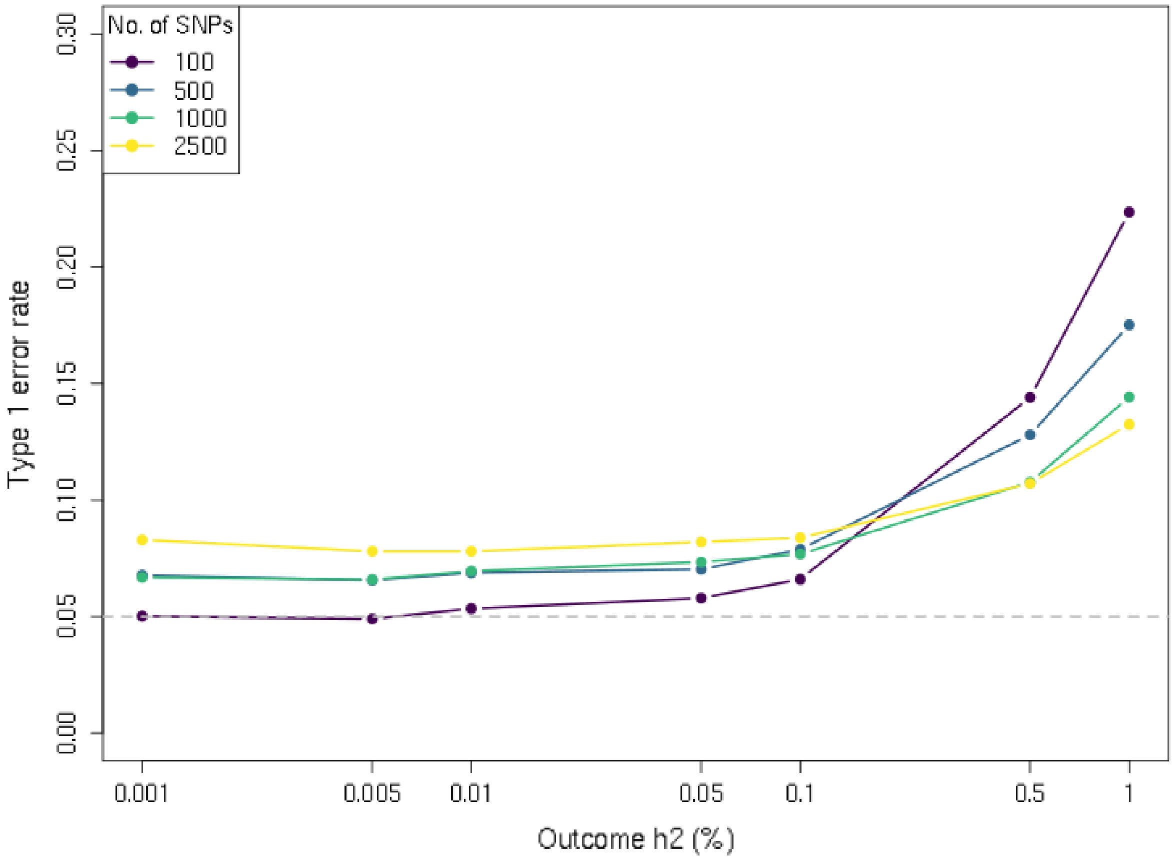


**Supplemental Figure 1. Type 1 error rate simulation results as a function of the number of SNPs in the gene.** *Shown is the type 1 error rate (at significance threshold of 0.05) of the LAVA-TWAS model for the null hypothesis of no genetic covariance ($cov(G_E, G_Y) = 0$), at different levels of local heritability (h2) for outcome phenotype (horizontal axis). Results are shown for sample sizes of 1,000 and 10,000 for the expression and outcome respectively, at 5% local heritability for the expression. As shown, the number of SNPs interacts with the local heritability of the outcome, with lower numbers of SNPs resulting in a lower type 1 error rate at lower heritability, but increasing more rapidly as a function of heritability than conditions with higher numbers of SNPs.*

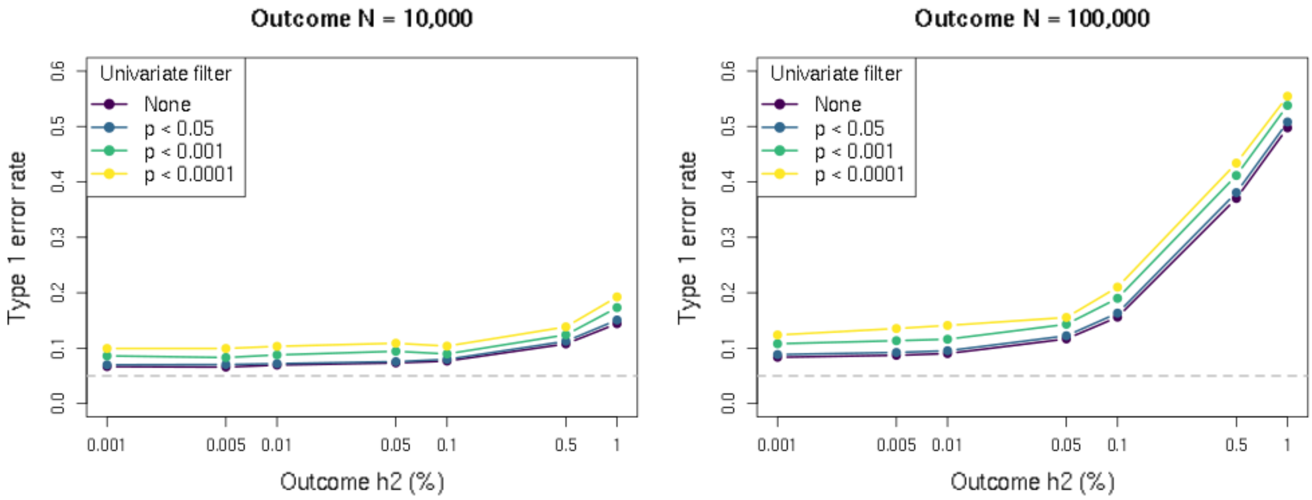


**Supplemental Figure 2. Type 1 error rate simulation results as a function of univariate input filtering.** *Shown is the type 1 error rate (at significance threshold of 0.05) of the LAVA-TWAS model for the null hypothesis of no genetic covariance ($cov(G_E, G_Y) = 0$), at different levels of local heritability (h2) for outcome phenotype (horizontal axis) and different gene filtering settings. The univariate filtering is based on the test of genetic signal for the gene expression implemented in LAVA, applying either no filtering or subsetting to iterations of the simulation with univariate p-value smaller than 0.05, 0.001 or 0.0001 (corresponding to multiple testing correction for 1, 50 or 500 genes respectively) before computing the type 1 error rate on the remaining iterations. Results are shown for a sample size (N) of 1,000 and a local heritability of 5% for the expression, and 1,000 SNPs. As shown, filtering at a univariate p-value threshold of 0.05 has little effect, but at the stricter thresholds the type 1 error rate becomes further increased.*

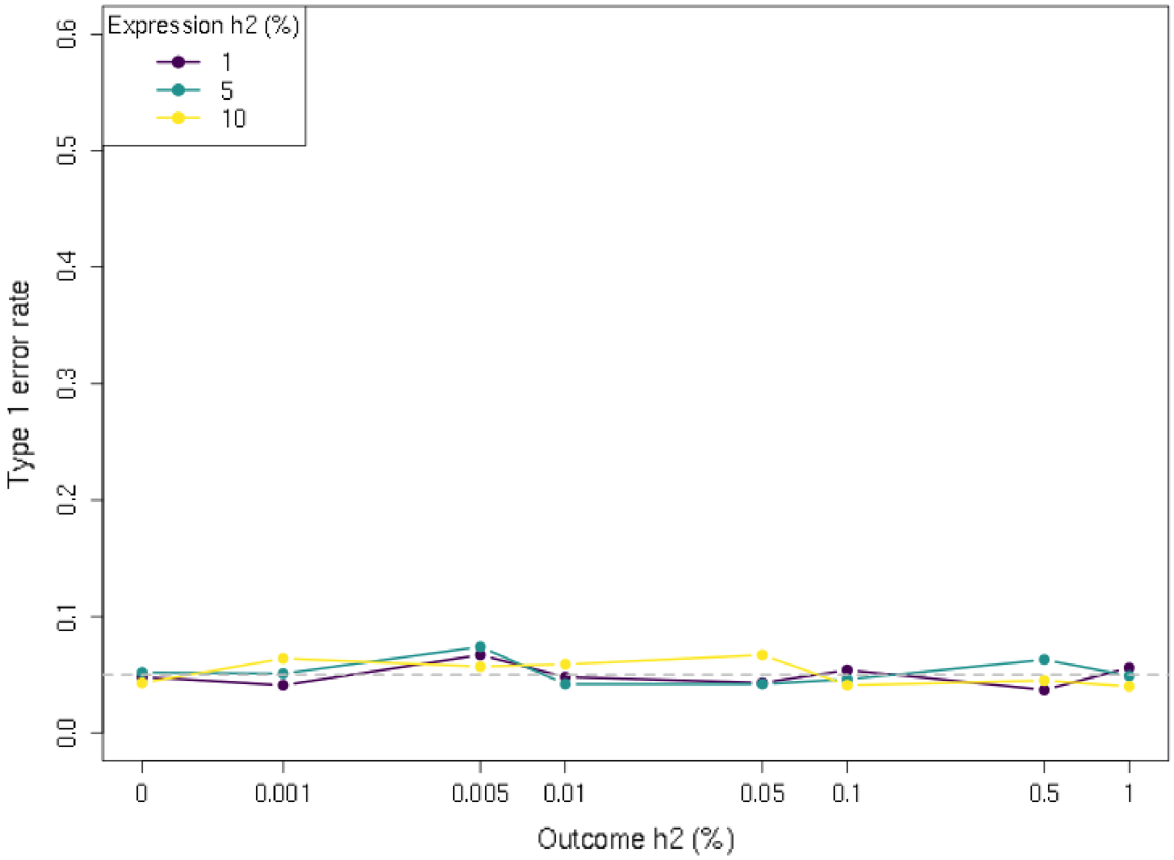


**Supplemental Figure 3. Type 1 error rate simulation results for LAVA-TWAS under the TWAS null hypothesis.** *Shown is the type 1 error rate (at significance threshold of 0.05) of LAVA-TWAS under the TWAS null hypothesis* $cov(\hat{G}_E, G_Y) = 0$*), at different levels of local heritability (h2) for outcome phenotype (horizontal axis) and gene expression (separate lines). Results are shown for sample sizes of 1,000 and 100,000 for the expression and outcome respectively, with 1,000 SNPs. As shown, and in contrast to the results when testing the null hypothesis of no genetic covariance (*$cov(G_E, G_Y) = 0$*, Figure 1), type 1 error rates here are well controlled regardless of condition.*

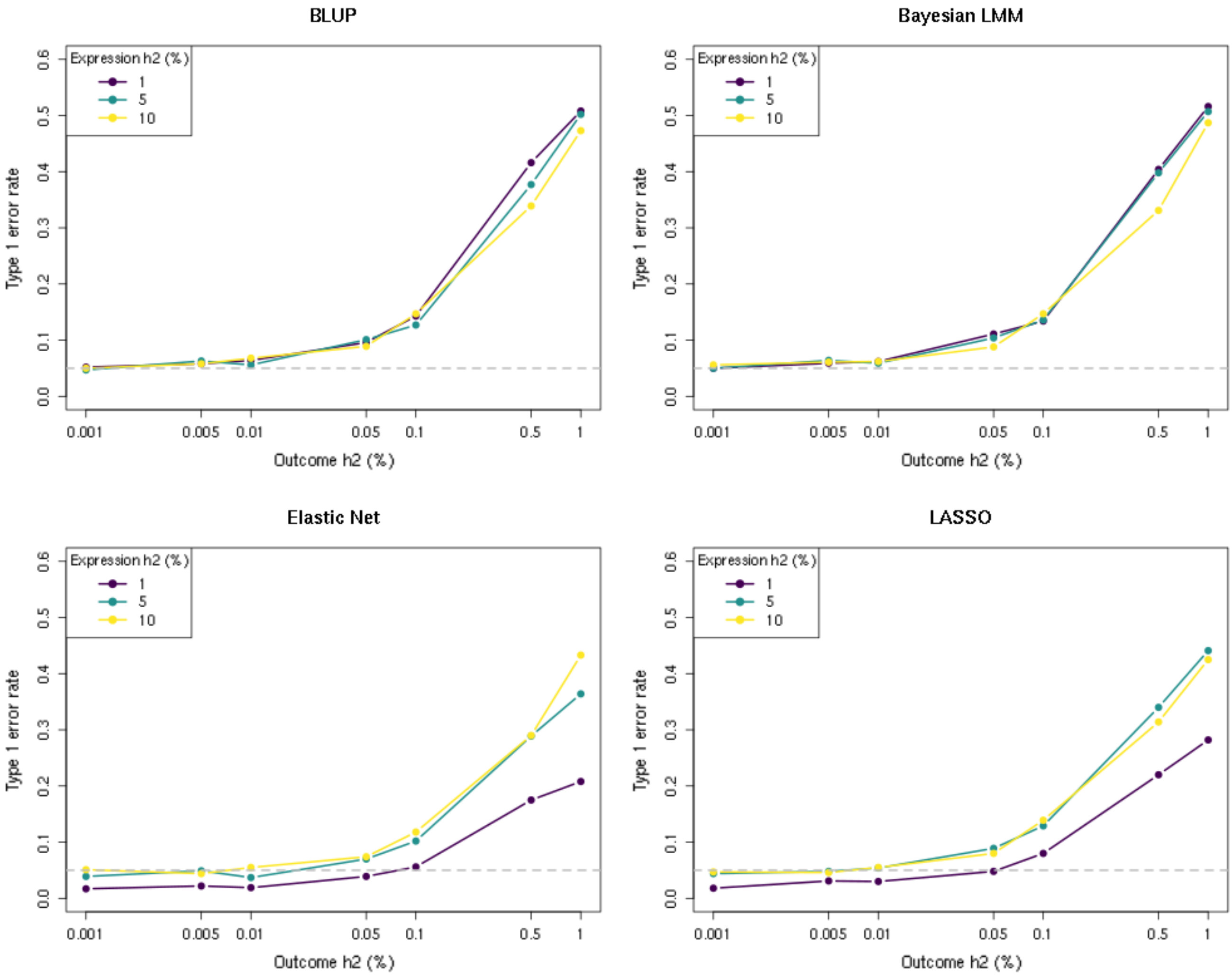


**Supplemental Figure 4. Type 1 error rate simulation results for FUSION models.** *Shown is the type 1 error rate (at significance threshold of 0.05) of the four penalized regression models in FUSION for the null hypothesis of no genetic covariance* ($cov(G_E, G_Y) = 0$)*, at different levels of local heritability (h2) for outcome phenotype (horizontal axis) and gene expression (separate lines). Results are shown for sample sizes of 1,000 and 100,000 for the expression and outcome respectively, with 1,000 SNPs, thus corresponding to the results for LAVA-TWAS shown in the bottom-right panel of Figure 1. As shown, the results are very similar across the different models. The Elastic Net and LASSO models do show decreased type 1 error rate for the 1% expression heritability condition, which is due to these models more frequently failing to converge if there is little detectable genetic signal for the expression, making the TWAS analysis inherently non-significant in those cases (whereas the BLUP and Bayesian LMM models always yield an* $\hat{\alpha}_E$*).*

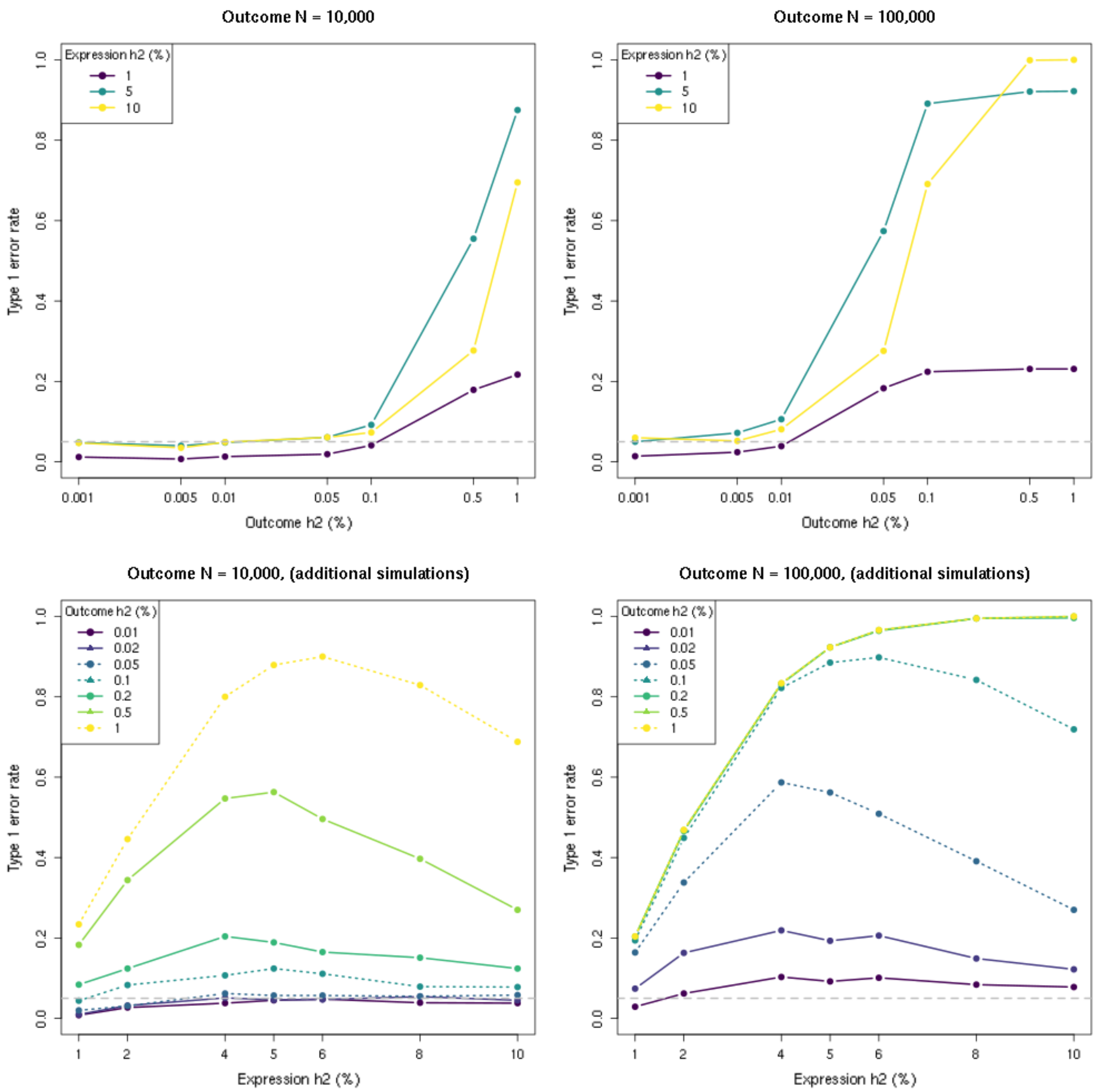


**Supplemental Figure 5. Type 1 error rate simulation results for CoMM.** *Shown on the top row is the type 1 error rate (at significance threshold of 0.05) of CoMM for the null hypothesis of no genetic covariance (*$cov(G_E, G_Y) = 0$*), at different levels of local heritability (h2) for outcome phenotype (horizontal axis) and gene expression (separate lines). Results are shown for sample sizes of 1,000 for the expression, and sample sizes of 10,000 (left) and 100,000 (right) for the outcome. Results in the top-right panel thus correspond to the LAVA-TWAS results in the bottom-right panel of Figure 1 and the FUSION results in Supplemental Figure 4, except for using 100 rather than 1,000 SNPs. As shown, type 1 error rate inflation is generally more pronounced than for the other methods. Moreover, there is a (partial) reversal of the effect of the local heritability of the expression, with the error rates for the 10% heritability level generally in between those of 1% and 5% heritability. Subsequently, additional simulations were performed using a more fine-grained range of heritability values to further investigate this phenomenon, with results shown on the bottom row with local heritability for the gene expression now on the horizontal axis and for the outcome as separate lines. These additional results confirm the non-monotonic effect of gene expression heritability on type 1 error rate, and demonstrate an interaction between it and the heritability and sample size of the outcome as well. See also section* Discussion of simulation results *above.*

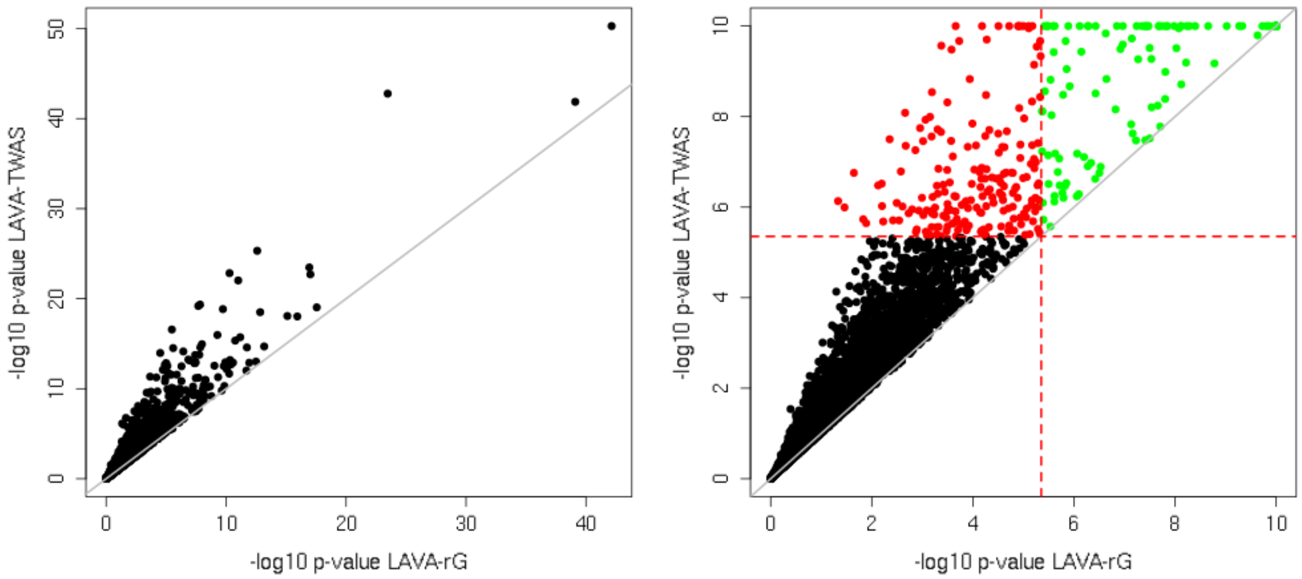


**Supplemental Figure 6. P-value bias in the real data analysis for BMI.** *Shown is the relation between the -log10 p-values of LAVA-TWAS model against those of the LAVA-rG model, demonstrating the bias in the LAVA-TWAS p-values as a result of its failure to account for the uncertainty in* $\hat{G}_E$. *On the left a scatterplot of all the -log10 p-values from the analysis, on the right the same values truncated to* $1.0 \times 10^{-10}$ *to better visualize the region around the Bonferroni-corrected significance threshold (horizontal and vertical dashed lines). In green the genes significant for both LAVA-TWAS and LAVA-rG, in red the invalid associations significant in LAVA-TWAS only.*

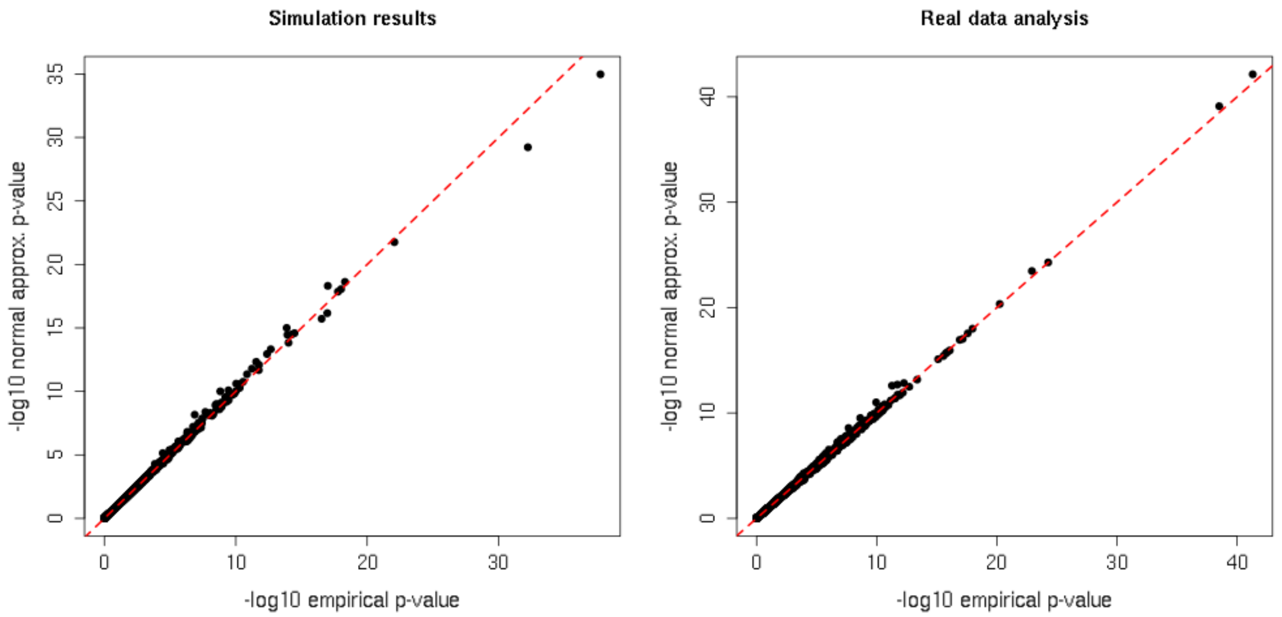


**Supplemental Figure 7. Comparison of -log10 p-values for different implementations of LAVA-rG.** *P-values in the published implementation of LAVA-rG are computed using a partially empirical, sampling-based approach. Shown here is a comparison with p-values computed using a normal approximation, results for which are used throughout this paper rather than those based on the empirical p-values. Here a comparison is given of all the LAVA-rG p-values from the simulations (left) and real-data analysis (right). As shown, these p-values are virtually identical, and as such results based on either approach can be considered interchangeable.*

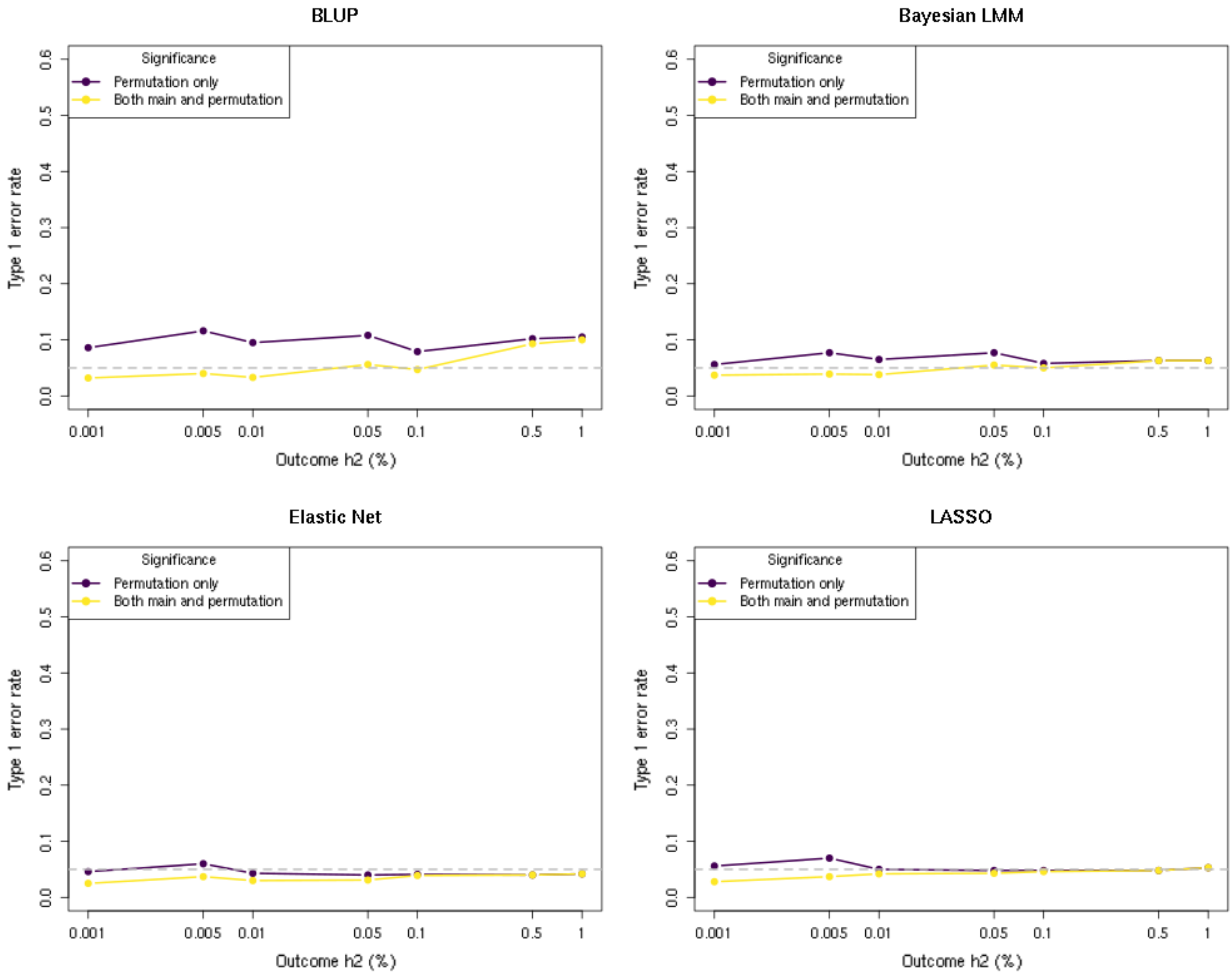


**Supplemental Figure 8. Type 1 error rate simulation results for FUSION models with post hoc permutation test.** *Shown is the type 1 error rate (at significance threshold of 0.05) of the four penalized regression models in FUSION for the null hypothesis of no genetic covariance ($cov(G_E, G_Y) = 0$), at different levels of local heritability (h2) for the outcome phenotype (horizontal axis). Results are shown for 5% heritability for the gene expression, and sample sizes of 1,000 and 100,000 for the expression and outcome respectively, with 1,000 SNPs. Significance is declared either when the permutation p-value is below 0.05 (black lines), or when the permutation p-value and the main analysis p-value are both below 0.05 (yellow lines). As shown, type 1 error rates are generally well controlled for the Elastic Net and LASSO models, and well controlled for the Bayesian LMM model when using both p-values, though for the BLUP model inflation remains even when using both p-values.*

### SUPPLEMENTAL TABLES

***Supplemental Table 1. Summary of results of TWAS and local genetic correlation analyses of published summary statistics for five phenotypes, with FUSION univariate filtering.***

| Phenotype | Sample size[a] | Number of SNPs[b] | Genes tested | Significance threshold | LAVA signif. associations $r_G$ | LAVA signif. associations TWAS | LAVA signif. associations Overlap %[c] | FUSION signif. associations Elast. net | FUSION signif. associations LASSO |
|---|---|---|---|---|---|---|---|---|---|
| Blood pressure[7] | 361K | 5.94M | 3748 | $1.33 \times 10^{-5}$ | 49 | 105 | 46.7 | 93 | 100 |
| BMI[8] | 807K | 6.28M | 3748 | $1.33 \times 10^{-5}$ | 98 | 273 | 35.9 | 291 | 313 |
| Type 2 diabetes[7] | 18.5K/366K | 5.94M | 3748 | $1.33 \times 10^{-5}$ | 20 | 54 | 37.0 | 28 | 33 |
| Educational attainment[9] | 766K | 6.18M | 3748 | $1.33 \times 10^{-5}$ | 40 | 134 | 29.9 | 141 | 147 |
| Schizophrenia[10] | 67.4K/94.0K | 6.08M | 3748 | $1.33 \times 10^{-5}$ | 37 | 121 | 30.6 | 122 | 127 |

Genes were included for testing if they exhibited significant univariate eQTL signal for FUSION at $\alpha_{BONF} = 0.05/14{,}584 = 3.43 \times 10^{-6}$. Since for FUSION the univariate p-values are computed in advance, before subsetting the data to SNPs overlapping with the GWAS input, the selected genes are the same for all phenotypes. Results were Bonferroni corrected per phenotype for the number genes tested. Note that all significant LAVA-rG associations were also significant in LAVA-TWAS.

[a] Showing case/control for binary phenotypes

[b] After filtering for overlap with 1,000 Genomes and GTEx SNPs

[c] Percentage of significant LAVA-TWAS associations that were also LAVA-rG associations

***Supplemental Table 2. Summary of results of TWAS and local genetic correlation analyses for genes with no detectable eQTL signal.***

| Phenotype | Sample size[a] | Number of SNPs[b] | Genes tested | Significance threshold | LAVA signif. associations $r_G$ | LAVA signif. associations TWAS | FUSION signif. associations Elast. net | FUSION signif. associations LASSO |
|---|---|---|---|---|---|---|---|---|
| Blood pressure[7] | 361K | 5.94M | 3962 | $1.26 \times 10^{-5}$ | 0 | 55 | 55 | 58 |
| BMI[8] | 807K | 6.28M | 5240 | $9.54 \times 10^{-6}$ | 2 | 205 | 205 | 254 |
| Type 2 diabetes[7] | 18.5K/366K | 5.94M | 3962 | $1.26 \times 10^{-5}$ | 0 | 38 | 17 | 20 |
| Educational attainment[9] | 766K | 6.18M | 5144 | $9.72 \times 10^{-6}$ | 0 | 86 | 90 | 96 |
| Schizophrenia[10] | 67.4K/94.0K | 6.08M | 4823 | $1.04 \times 10^{-5}$ | 0 | 77 | 60 | 82 |

Genes were included for testing if they exhibited no univariate eQTL signal in the LAVA univariate test, with univariate p-value > 0.05. Results were Bonferroni corrected per phenotype for the number genes tested.

[a] Showing case/control for binary phenotypes

[b] After filtering for overlap with 1,000 Genomes and GTEx SNPs

***Supplemental Table 3. Summary of results of FUSION permutation tests for published summary statistics for five phenotypes.***

| Phenotype | LAVA signif. associations $r_G$ | LAVA signif. associations TWAS | FUSION Elastic Net signif. Associations Main | FUSION Elastic Net signif. Associations Permutation | FUSION Elastic Net signif. Associations Both | FUSION LASSO signif. Associations Main | FUSION LASSO signif. Associations Permutation | FUSION LASSO signif. Associations Both |
|---|---|---|---|---|---|---|---|---|
| Blood pressure[7] | 68 | 107 | 100 | 1 | 1 | 109 | 2 | 1 |
| BMI[8] | 134 | 232 | 297 | 3 | 0 | 328 | 11 | 1 |
| Type 2 diabetes[7] | 27 | 45 | 26 | 0 | 0 | 33 | 13 | 0 |
| Educational attainment[9] | 59 | 108 | 149 | 2 | 0 | 165 | 1 | 0 |
| Schizophrenia[10] | 53 | 92 | 109 | 1 | 0 | 114 | 3 | 1 |

Genes were included for bivariate testing if they exhibited significant univariate eQTL signal in the LAVA univariate test at $\alpha_{BONF} = 0.05/14{,}584 = 3.43 \times 10^{-6}$. Results were Bonferroni corrected per phenotype for the number genes tested, see Table 2 for the number of genes tested and resulting significance threshold. FUSION permutation test used an adaptive permutation procedure with up 10,000,000 permutations.
